# Age-related changes in behavioural and neural variability in a decision-making task

**DOI:** 10.1101/2025.08.22.671763

**Authors:** Fenying Zang, Anup Khanal, Sonja Förster, International Brain Laboratory, Anne K. Churchland, Anne E. Urai

## Abstract

Age-related cognitive decline in learning and decision-making may arise from increased variability of neural responses. Here, we investigated how ageing affects behavioural and neural variability by recording >18,000 neurons across 16 brain regions (including cortex, hippocampus, thalamus, midbrain, and basal ganglia) in younger and older mice performing a visual decision-making task. Older mice showed more variable response times, reproducing a common finding in human ageing studies. Ageing globally increased firing rates and post-stimulus neural variability (quantified using the Fano factor), and decreased variability quenching–the reduction in neural variability upon stimulus presentation. Older animals showed higher overall firing rates across areas of visual and motor cortex, striatum, midbrain, and hippocampus, but lower firing rates in thalamic areas. Age-related attenuation in stimulus-induced variability quenching was most prominent in visual and motor cortex, striatum, and thalamic area. These findings show how large-scale neural recordings can help uncover regional specificity of ageing effects in single neurons, ultimately improving our understanding of the neural basis of age-related cognitive decline.

## Introduction

Age-related cognitive impairments have long been thought to arise from higher levels of neural noise or variability. In the second half of the 20th century, theoretical accounts proposed that increased age-related neural variability reduces the effective signal-to-noise ratio within the central nervous system, leading to cognitive impairments^1–4^. Behavioural studies at the time supported this idea, showing that the absolute threshold for detecting stimuli increases with age^4^ and that adding random stimulus noise (simulating internal neural noise) could mimic age-related differences in task performance, particularly in visual tasks^1^. More recent theoretical accounts propose that declining neuromodulation in healthy ageing may impair neural networks’ gain control, the efficient modulation of a neuron’s input-output function^5^. Such changes reduce the precision of neural representations, implying that neural dedifferentiation lies at the basis of cognitive changes in healthy ageing^6^.

Human studies of healthy ageing, typically comparing young adults (20–30 years) with older adults (often up to ∼85 years), present a complex and sometimes contradictory picture of age-related changes in neural variability. While some studies found widespread age-related increases in blood oxygen level-dependent (BOLD) variability across both cortical and subcortical regions^7^, others reported that many brain regions become less variable in older people^8–10^. The behavioural implications of age-related changes in BOLD variability remain debated. Some studies find lower brain variability in older people, linked to slower and more variable response times^9^, while others report increased BOLD variability in older adults, negatively affecting decision-making^11^. Older adults’ EEG activity consistently shows an increase in neural variability, as measured by metrics such as P300 latency variability^12^, weighted permutation entropy^13^, and particularly the slope of 1/f power spectra^14–16^. The latter measure, characterized by the aperiodic component of 1/f power spectral density^16^, has been interpreted as reflecting an increase in baseline neural noise, and correlates with age-related impairments in cognitive functions such as visual working memory^16^ and visual processing^15^. However, flatter slopes of EEG power spectra were recently shown to arise from a cardiac, rather than neural, source that changes with age^17^. This highlights the importance of using direct neural recordings to study brain function without age-related vascular and cardiac confounds that may affect the interpretation of BOLD or EEG data^17^.

Observations from single-neuron measurements in cortical areas show a complex picture of age-related change. Some studies report increased firing rates^18–21^, while others have found decreased rates with ageing^22^. Only a few studies have looked directly into age-related changes in the variability of single-neuron activity. In anaesthetized rhesus monkeys, comparing young adults (5–9 years; ∼18–30 human-equivalent years) with aged animals (23–31 years; ∼70–90 human-equivalent years), cortical areas V1 and MT show increased age-related trial-by-trial spiking variability (measured using Fano factor, defined as the ratio of spike-count variance to mean spike count across trials), reduced signal-to-noise ratio^21^, and increased noise correlations^23^. Extending this evidence to rodents, a recent in vivo study in mouse V1 reported higher single-neuron Fano factor and increased noise correlations in 12-month-old mice compared with 8-week-old mice in response to repeated grating stimuli^24^. Although informative, these findings are limited by relatively small datasets: for example, Yang et al. recorded 172 and 173 neurons from 3 young and 4 old monkeys, respectively^21^, while the recent mouse V1 study recorded 136 neurons from 6 young mice and 140 neurons from 5 old mice^24^. Moreover, their generalization to task-engaged animals and other brain regions remains unclear. In addition, these studies did not examine the time course of neural variability. Neural variability, as measured by the Fano factor, typically decreases following stimulus onset—a phenomenon known as variability quenching—which has been consistently observed across species and cortical areas^25–29^. Yet, how this reduction of variability in response to sensory stimuli changes with ageing remains unknown, particularly in behaving animals.

More broadly, rodent studies have also reported age-related changes in neural activity across multiple levels of organization, from cellular and synaptic changes in excitability, synaptic transmission, and excitation–inhibition balance to circuit- and population-level alterations in firing rates and population activity patterns across regions^30–37^. Related changes have also been reported in rodent models of neurodegenerative disease, including Alzheimer’s disease, which are often characterized by circuit hyperexcitability^38–40^. These studies span diverse experimental contexts, but often focus on one or a small number of regions and include many experiments conducted in passive paradigms or in anesthetized/ex vivo preparations. As a result, they provide limited insight on how ageing impacts trial-to-trial neural variability during perceptual decision-making across brain regions.

In recent years, large-scale neural recording techniques have offered exciting opportunities to record from many neurons as animals perform complex decision-making tasks^41^. We here build on the work of the International Brain Laboratory (IBL), which has standardized training and recording pipelines for mouse decision-making behaviour^42–44^. Recording brain-wide neural activity during a visual decision-making task revealed that decision-related signals are distributed across much of the mouse brain^45^. Visual stimuli evoked transient responses in early visual areas, which were followed by choice-related ramping in midbrain and hindbrain regions. Neural signals related to movement, reward, and feedback were widespread across the brain^45^. Here, we build on this well-curated dataset and contribute additional recordings from older mice, allowing us to investigate age-related changes in behaviour and neural activity. We openly share all data and code to promote reproducibility and further analyses of this rich dataset.

In this study, we use large-scale neural recordings from behaving mice to investigate age-related changes in single-neuron variability across the brain. We analysed extracellular Neuropixels recordings from 149 mice (aged 3–20 months; roughly corresponding to 23–70-year-old humans^46^) across 16 brain regions (in cortex, striatum, midbrain, hippocampus, and thalamus; Table 1) while animals performed a standardized perceptual decision-making task ^43^.

**Table 1.**
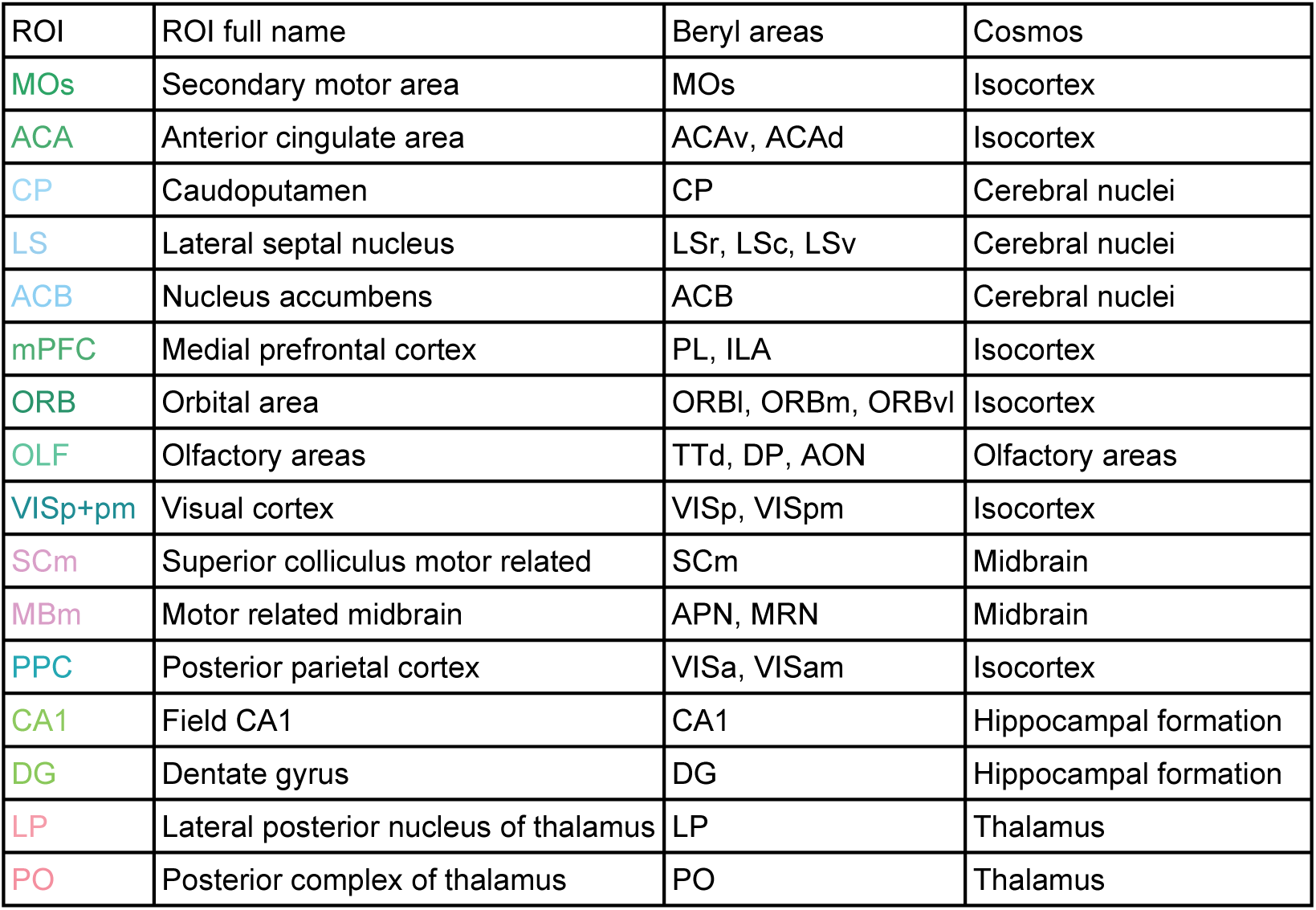
ROI definitions. This table details the composition of each defined ROI, including its abbreviation and the constituent brain regions.

| ROI | ROI full name | Beryl areas | Cosmos |
| --- | --- | --- | --- |
| MOs | Secondary motor area | MOs | Isocortex |
| ACA | Anterior cingulate area | ACAv, ACAd | Isocortex |
| CP | Caudoputamen | CP | Cerebral nuclei |
| LS | Lateral septal nucleus | LSr, LSc, LSv | Cerebral nuclei |
| ACB | Nucleus accumbens | ACB | Cerebral nuclei |
| mPFC | Medial prefrontal cortex | PL, ILA | Isocortex |
| ORB | Orbital area | ORBI, ORBm, ORBvl | Isocortex |
| OLF | Olfactory areas | TTd, DP, AON | Olfactory areas |
| VISp+pm | Visual cortex | VISp, VISpm | Isocortex |
| SCm | Superior colliculus motor related | SCm | Midbrain |
| MBm | Motor related midbrain | APN, MRN | Midbrain |
| PPC | Posterior parietal cortex | VISa, VISam | Isocortex |
| CA1 | Field CA1 | CA1 | Hippocampal formation |
| DG | Dentate gyrus | DG | Hippocampal formation |
| LP | Lateral posterior nucleus of thalamus | LP | Thalamus |
| PO | Posterior complex of thalamus | PO | Thalamus |

We found that older animals showed higher trial-to-trial variability in response times. Neural recordings show that ageing is accompanied by higher firing rates across the brain, increased post-stimulus neural variability (as measured with Fano factors), and attenuated stimulus-induced variability quenching. Different metrics to quantify age-related neural changes show complex regional patterns. These results suggest that behavioural differences in older animals may be accompanied by specific changes in neural variability after stimulus onset.

## Results

We combined a previously released dataset of extracellular Neuropixels recordings^47^ from young mice (N = 130, 89 male, mean age = 6.64 months, range 3.10–15.13 months)^45^ with additional neural recordings in older mice (N = 19, 11 male, mean age = 16.58 months, range 10.58–19.90 months), all acquired using standardized protocols for behaviour and neural recordings (Figure 1a; see Methods)^43,45^. Briefly, mice were trained to discriminate the location of a visual stimulus on the left or right side of the screen by turning a small steering wheel in front of them. The stimuli varied in contrast, modulating task difficulty. After applying a rigorous set of standardized quality control measures (see Methods), we included a total of 149 mice that performed 367 sessions and underwent 503 Neuropixels insertions. Animals’ age at the day of recording ranged from 3–20 months (Figure 1b), roughly corresponding to a human age range of 23–70 years^46^—approximately spanning the typical working lifespan from early adulthood to retirement. For visualization, mice were categorized based on their age at recording into a young group (N = 97, mean age = 5.60 months) and an old group (N = 52, mean age = 12.07 months), using 7.6 months (mean age in the dataset) as the age cutoff (Figure 1b). Age was treated as a continuous variable in all statistical analyses.

**Figure 1.**
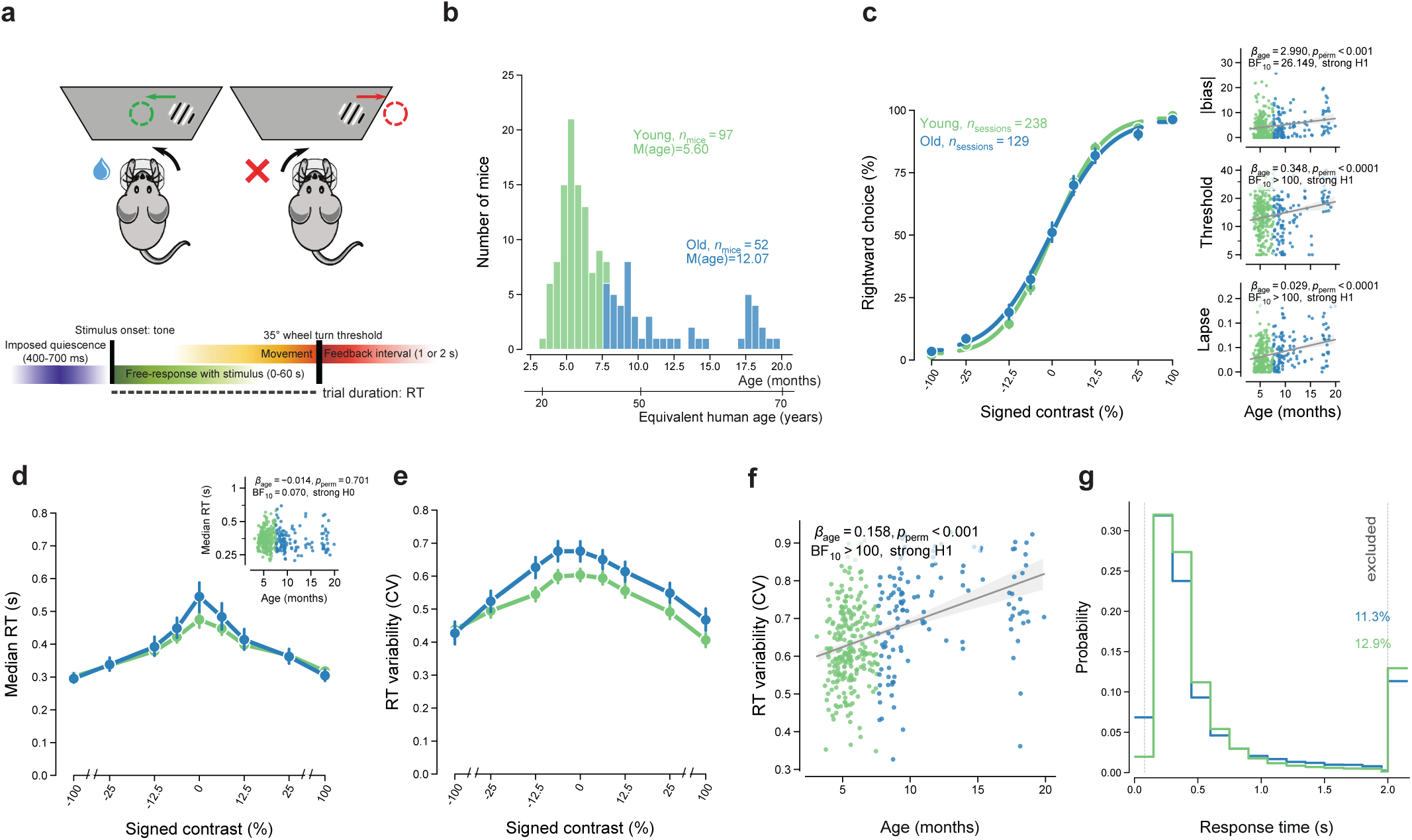
Older mice show larger RT variability and slightly worse performance in a standardized decision-making task. (a) Schematic of the task, showing a correct response (the visual stimulus was brought to the centre of the screen) vs. a wrong response (the visual stimulus was moved off the screen). Response times (RT) were defined as the interval from stimulus onset to response completion. Adapted from ref. 45 (<u>International Brain Laboratory et al.</u>; © The Author(s) 2025; licensed under <u>CC BY 4.0</u>). (b) Distribution of mouse age on the day of neural recording (n = 149 mice; young n = 97; old n = 52; age cutoff: 7.6 months). Group splits were only used for visualization; all statistics used age as a continuous variable. The age range roughly corresponds to a human age range of 23–70 years^46^. (c–f) Behavioural statistics were computed at the session level (n = 367 sessions; young n = 238; old n = 129). Age effects were estimated using general linear models with age as a continuous predictor. All statistical conclusions were based on Bayes Factors (BF_10_); two-sided permutation p-values are unadjusted for multiple comparisons and are reported as complementary statistics (see Methods). Error bars and shaded bands indicate 95% confidence intervals. Unless otherwise noted, regression lines are plotted only when the Bayes Factor indicates strong or moderate H1. (c) Left: average psychometric curve across sessions for each age group. Right: relationship between psychometric parameters and mouse age. Each dot represents one session. Absolute bias: *p*perm =0.00069; threshold: *p*perm =0.00009; lapse: *p*perm =0.00009. Note the log-scaling on the y-axis of the threshold panel. (d) Average chronometric curve across sessions for each age group. Inset: Relationship between median RT and mouse age. Median RT was calculated across all included trials, irrespective of stimulus contrast. *P*perm =0.70093. Note the log-scaling on the y-axis. (e) RT variability, quantified by coefficient of variation (CV), across stimulus conditions for each age group. (f) Relationship between RT variability and mouse age. *p*perm =0.00009. (g) RT distribution by age group. Bars at the far left and far right show the percentage of trials excluded due to RT shorter than 0.08 s or longer than 2 s (see also Supplementary Figure 3).

### Older mice show more variable behaviour on a standardized decision-making task

Older animals did not take longer to learn the standardized task, and animals of different ages eventually met the standardized criteria to be classified as trained (Supplementary Figure 1). Although animals that took longer to learn were older by the time of neural recording, the main source of age variability at recording time arises from starting the training program later (Supplementary Figure 2). Only behavioural data from the recording sessions were included in our main analyses.

During recording sessions, older mice performed slightly more trials until automated behaviour-based stopping criteria ended the session (range: 401–1279; Supplementary Figure 3a)^43^. This may reflect larger weight-dependent water requirements in older, heavier animals. To control for this potential confound in our statistical analyses (especially when computing neural metrics sensitive to trial counts), we restricted our analyses to the first 400 trials of each session. Additionally, only trials with RTs between 80 ms and 2 s were included, to exclude anticipatory (<80 ms) and very slow (>2 s) responses that are unlikely to reflect stimulus-driven decisions (Figure 1g), consistent with prior IBL analyses^45^. Unless otherwise specified, regression lines are shown only when the Bayes Factor indicates strong or moderate evidence in favour of H1. The number of in- or excluded trials did not show a significant correlation with age (included trial counts: Supplementary Figure 3c; number of trials excluded by RT filtering: Supplementary Figure 3b).

Older mice performed slightly worse on the task. Psychometric curves fit to each session (Figure 1c) showed that three psychometric function parameters—absolute bias, threshold, and mean lapse rate—showed an increase with age (Figure 1c, right panels). We did not observe age-related changes in choice bias conditioned on blocks or previous choices (Supplementary Figure 4), suggesting that the age-related differences in overall psychometric parameters are not driven by changes in the integration of prior information with sensory evidence. While history-dependent biases are well established in this task^48^, they do not seem related to the age effects observed here.

Older mice did not respond more slowly, but were more variable in their response times (RTs). We defined RT as the time animals took to complete their choice (distinct from the time at which they initiated their first movement). Although older mice did not exhibit slower RTs on average (Figure 1d, right), their RTs were notably more variable (Figure 1e, f). The variability in RTs, as measured by the coefficient of variation (CV; standard deviation / mean), increased with age. This finding aligns with many human studies that demonstrate increased trial-to-trial RT variability in older adults across a wide range of cognitive tasks^5,49–51^. These results suggest that age-related increases in trial-to-trial RT variability, widely reported in humans, can also be observed in behaving rodents. This result was robust to various control analyses: other measures of variability (Supplementary Figure 5a) and for a definition of RT that captures the first movement initiation, rather than movement completion (Supplementary Figure 5b), and differences between dataset sources or lab-specific environmental factors (Supplementary Figure 6 and 7).

### Large-scale Neuropixels recordings across the mouse lifespan

To investigate age-related changes in single-neuron activity, we analysed data from 149 mice, comprising 367 sessions and 503 insertions (up to two per recording session) of Neuropixels recordings across 16 brain regions of interest (Figure 2a,b; Table 1). These included 6 cortical regions, along with structures from the hippocampus, thalamus, midbrain, basal ganglia, and olfactory areas (Figure 2c). These regions were selected based on a combination of scientific, statistical, and practical considerations: scientifically, frontal cortical areas are known to show early age-related decline; statistically, the set includes the repeated site used to assess cross-laboratory reproducibility^44^; and practically, these areas offered reliable surgical accessibility. We applied a series of standardized quality control metrics to the neural data before proceeding with subsequent analyses (Figure 2d; see Methods for details)^45^. From 242,671 recorded units (including multi-neuron activity) across these 16 regions, the quality control process identified 18,755 good neurons. Overall neural yield was slightly reduced in older mice (*β*age=-0.106, *p*perm=0.001; BF_10_ >100, strong H1), specifically in PPC, LS and ACA (Supplementary Figure 8).

**Figure 2.**
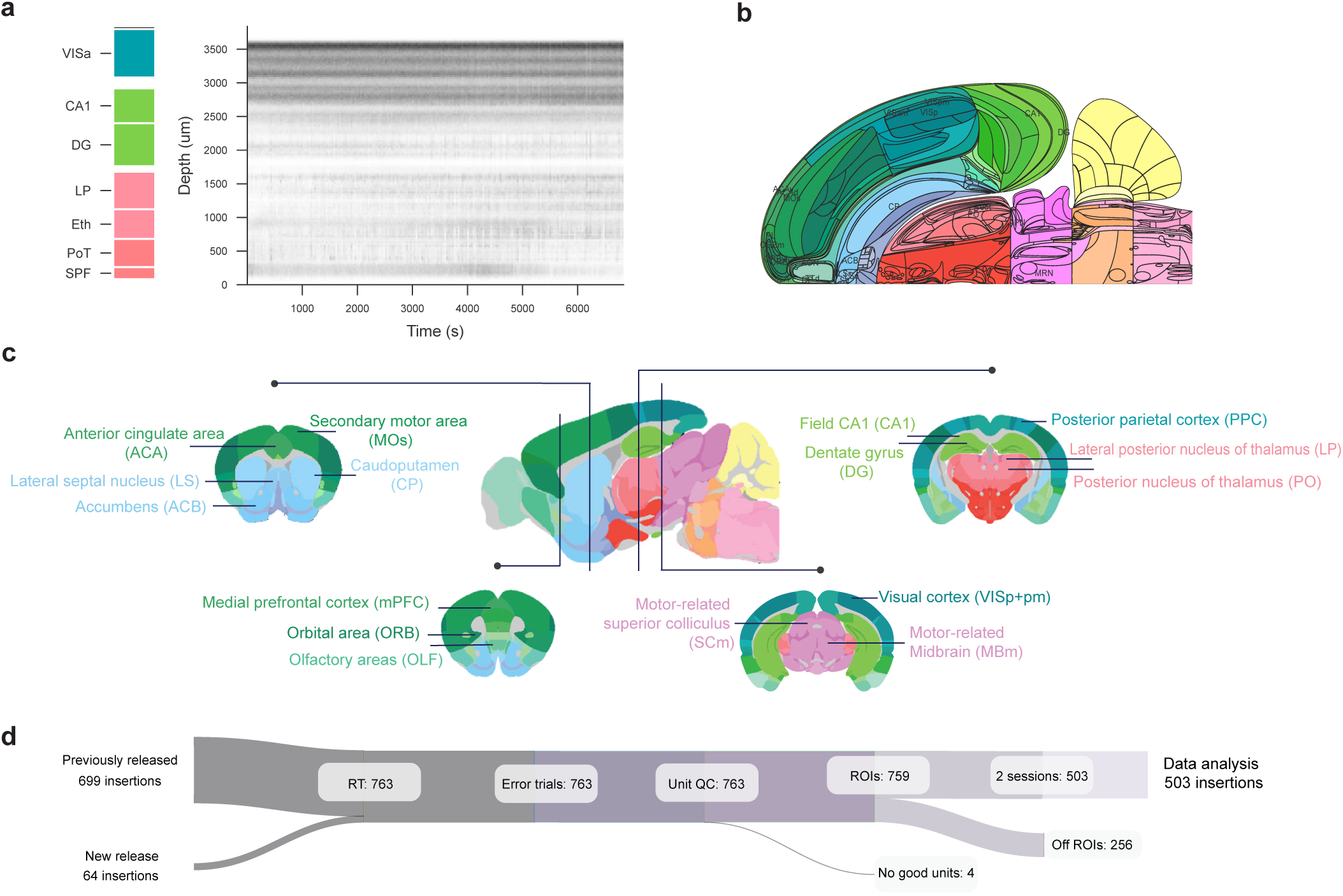
Large-scale neural recording across the mouse brain. (a) Raster plot from one example recording probe, with aligned regions indicated on the left. Brain regions: VISa (anterior visual area), CA1 (field CA1), DG (dentate gyrus), LP (lateral posterior nucleus of the thalamus), Eth (ethmoid nucleus of the thalamus), PoT (posterior triangular thalamic nucleus), SPF (subparafascicular nucleus). The y-axis shows brain regions this probe traverses, and the x-axis shows the time elapsed from session start. Each dot indicates one spike, with dark bands showing brain regions with many spiking neurons. (b) The 2D flatmap representation of the mouse brain^45^. Labelled regions indicate regions of interest (ROIs), as detailed in (c) and Table 1. (c) A 2D-sagittal mouse brain slice and four corresponding coronal slices, adapted from the Allen Mouse Brain Atlas and Allen Reference Atlas – Mouse Brain^102,103^ (mouse.brain-map.org and atlas.brain-map.org), with ROI acronyms and annotations added. (d) The number of insertions remaining after each quality control (QC) step. Previously released: the public dataset from IBL; new release: dataset recorded during the present project. RT: RT and missing events; Error trials: minimum 3 error trials; Unit QC: single unit QC; ROIs: ROIs only; 2 sessions: minimum 2 sessions per region. See Table 2 and Methods for more details.

**Table 2.** Quality control process. The table shows the step-by-step filtering of sessions and insertions, using the inclusion criteria described in the main text. The numbers in the first row are based on the data release.

| QC | level | Previously released dataset<sup>45</sup> |  | Newly recorded dataset |  |
| --- | --- | --- | --- | --- | --- |
|  |  | n_sessions | n_insertions | n_sessions | n_insertions |
| Session and insertion QC | insertion, session | 459 | 699 | 38 | 64 |
| Response time and missing events | trials | 459 | 699 | 38 | 64 |
| Minimum 3 error trials | session | 459 | 699 | 38 | 64 |
| Single unit QC | neuron | 458 | 695 | 38 | 64 |
| ROIs only | neuron | 329 | 439 | 38 | 64 |
| Minimum 2 sessions per region | neuron | 329 | 439 | 38 | 64 |

After confirming high-quality neural data across age groups, we first assessed age-related changes in two general measures of neural functioning: overall firing rates and contrast modulation of neural responses. Throughout, we present different neural metrics first globally across recorded brain regions and then show their regional distribution.

### Increased firing rate in older animals

Ageing increased firing rates globally. Previous studies have reported increased firing rates with ageing across species. In rodents, age-related increases in firing rates have been reported in multiple brain areas, including sensory cortex and hippocampus^33,52,53^. Increased firing rates have also been reported in the sensory cortex of other mammals, including cat V1^19^. In non-human primates, ageing has been associated with increased firing rates in cortical areas such as macaque V1 and MT^20,21^, as well as in specific cell types in the monkey prefrontal cortex^18^. In contrast, other studies have reported decreased firing rates with ageing, particularly in the monkey prefrontal cortex^22^. Here, when pooling neurons across all brain regions, we found global age-related changes in firing rates in both pre- and post-stimulus time windows (Figure 3a; Figure 3b,c, left panels). These effects were present in areas of visual and motor cortex (VISp+pm, MOs), striatum (LS, ACB, CP), midbrain (SCm, MBm), and hippocampus (CA1, DG) (right panels of Figure 3b,c; Supplementary Figure 9 and 10). Notably, the two thalamic areas LP and PO showed the reverse pattern of reduced firing rates in older animals (right panels of Figure 3b,c; Supplementary Figure 9 and 10). In PPC, an area causally implicated in age-related effects on learning and decision making^54^, we found evidence for no effect of ageing on firing rates. Frontal cortical, orbital, and cingulate areas showed weak or inconclusive ageing patterns.

**Figure 3.**
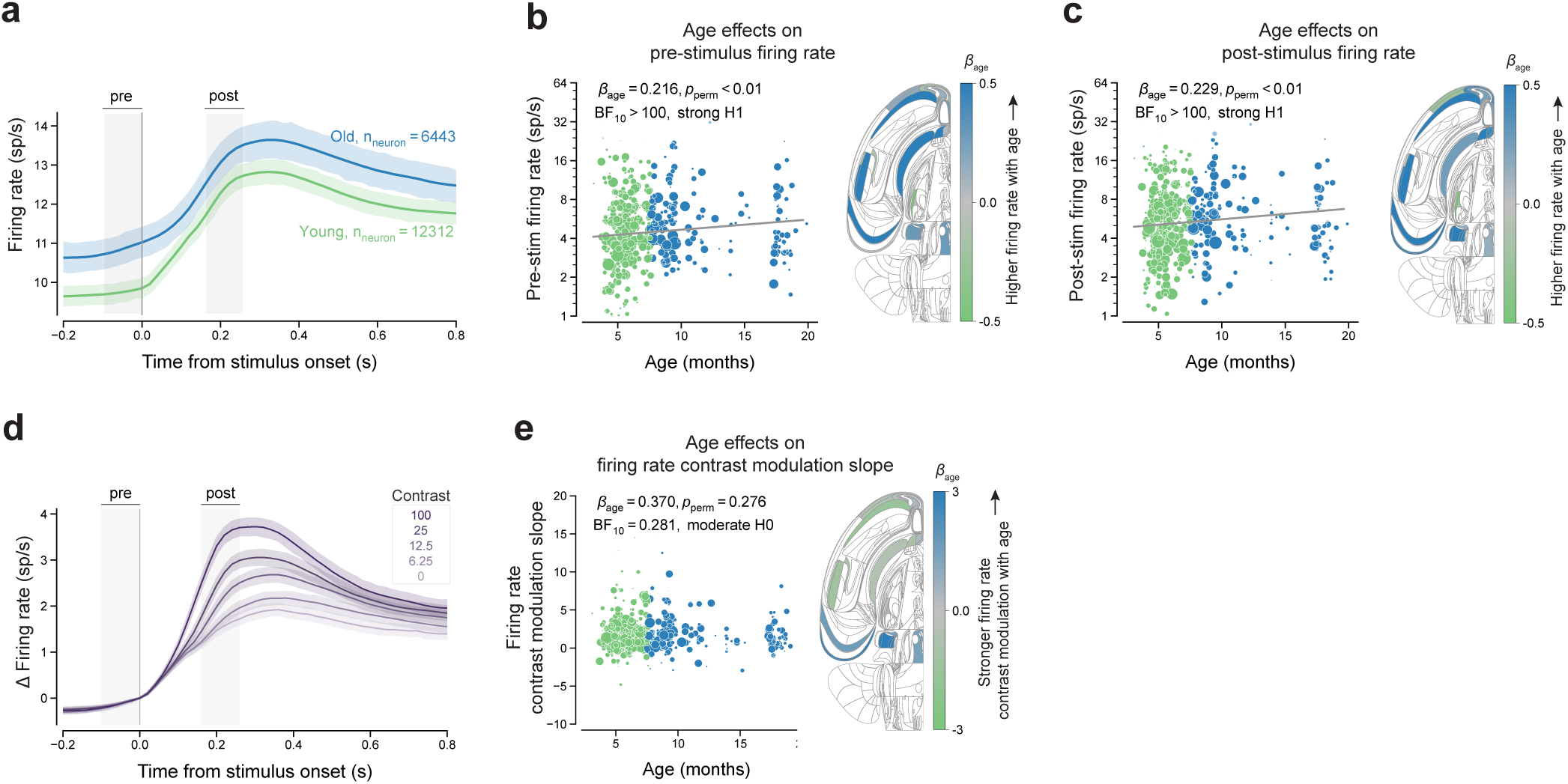
Age effects on global firing rates and neural contrast modulation. (a) Time courses of overall firing rates, aligned to stimulus onset. Thick lines represent the mean firing rate for each age group; shaded areas show 95% confidence intervals obtained by pooling neurons within each age group. The analysis included n = 18,755 neurons from n = 503 insertions and n = 149 mice. Sliding window width, 100 ms; step size, 20 ms. Grey areas indicate the pre-stimulus (-100 to 0 ms) and post-stimulus (160 to 260 ms) windows. For left panels in b, c and e, dots show insertion-level summaries for visualization, with dot size indicating the number of neurons in each insertion. Statistical analyses and fitted regression lines were based on single-neuron metrics using age as a continuous predictor. Age effects were estimated as *β*age using general linear models with age as a continuous predictor. All statistical conclusions were based on Bayes Factors ( BF_10_); two-sided permutation p-values are unadjusted for multiple comparisons and are reported as complementary statistics (see Methods). (b) Left: relationship between pre-stimulus firing rate and mouse age. Note the log-scaled y-axis. *p*perm=0.00899. Right: a flatmap representation showing region-specific age effects on pre-stimulus firing rate. The colour bar indicates the slope of age. White areas were not covered in our recordings. (c) Same as (b), but for post-stimulus firing rate. *p*perm=0.00200. (d) Average firing rate time courses across contrast levels. Saturation represents different contrast levels; shaded areas indicate 95% confidence intervals. (e) Same as (b), but for contrast modulation slope. Contrast modulation slopes were computed for each neuron by estimating the slope between different stimulus contrast levels and the change in baseline-corrected firing rates. *p*perm=0.27572. Because the omnibus analysis does not support a global age effect, this flatmap representation is shown for completeness and is not interpreted in detail in the main text.

Neural responses throughout the brain were modulated by stimulus contrast^45^ (Figure 3d, Supplementary Figure 11), but we found no significant effects of ageing on this global contrast modulation (Figure 3e, left panel). Previous studies show that aged neurons in the primary visual cortex of both cats and macaques exhibit weaker tuning to specific features like orientation and direction, and lower signal-to-noise ratios^19,20^, and aged mice show declines in frequency selectivity within the auditory midbrain^55^. This neural dedifferentiation, characterized by a loss of selectivity, means individual neurons respond to a broader range of stimuli, potentially leading to a less precise and noisier representation of sensory information^6^. We computed a contrast modulation slope for each neuron by fitting a linear regression between stimulus contrast levels [0, 6.25, 12.5, 25, 50, 100%] and the change in baseline-corrected firing rate (Δ firing rate, defined as post-stimulus minus pre-stimulus firing rate). An omnibus test revealed no significant age effects on contrast modulation overall (Figure 3e, left panel). Region-wise estimates are shown on a flatmap representation of the brain^45^ (Figure 3e, right panel) for completeness, but we do not interpret these regional patterns further given the lack of a robust omnibus effect. Notably, while *β*age estimates showed heterogeneous and bidirectional regional trends (Figure 3e, right panel; Supplementary Figure 12), we describe regional patterns in detail only when a corresponding global age effect is supported, in order to avoid over-interpretation of region-specific effects.

### Reduced stimulus-induced variability quenching in older animals

To quantify single-neuron response variability, we next computed Fano factors. Neural responses vary substantially across trials with identical external stimuli. The Fano factor, defined as the ratio of spike-count variance to mean spike count, serves as a measure of this single-neuron spiking variability, with its conceptual roots extending back to early analyses of neural firing patterns^56–59^. While in vivo spike trains often approximate Poisson statistics (characterized by a Fano factor of 1, indicating that the variance of spike counts equals the mean, Supplementary Figure 13), studies using Fano factor have revealed significant regional differences in spiking variability across brain regions^29,60–62^, as well as phenomena like quenching, where variability decreases at stimulus onset^25^. Reporting neural variability as Fano factor also facilitates direct comparison with prior work in this line of research^21,23,24^. To control for biases in Fano factor estimates due to condition-specific differences in mean firing rates, we applied a mean-subtraction correction method (see Methods)^63^.

Time courses showed different effects of ageing on the Fano factor between pre- and post-stimulus periods (Figure 4a). Ageing did not affect pre-stimulus Fano factors (Figure 4b, left panel), suggesting that baseline neural variability remains stable with age. Following stimulus presentation, however, older animals exhibited increased Fano factors (Figure 4c, left panel). This effect was most robustly observed in the midbrain (SCm, MBm) and thalamic area PO (Figure 4c, right panel; Supplementary Figure 14c). In contrast, cortical, striatal, and hippocampal areas showed only weak or inconclusive evidence of a post-stimulus ageing effect on Fano factor, showing a regionally specific pattern of age-related neural variability. These findings are consistent with previous work in anesthetized rhesus monkeys that reported increased Fano factors in V1 and MT neurons of older animals^21^. Our findings extend these observations by demonstrating that this age-related increase in neural variability is not a property of baseline activity but follows stimulus onset. We also note a short-lived transient Fano factor increase immediately after stimulus onset (more pronounced in older mice), the origin of which remains to be determined.

**Figure 4.**
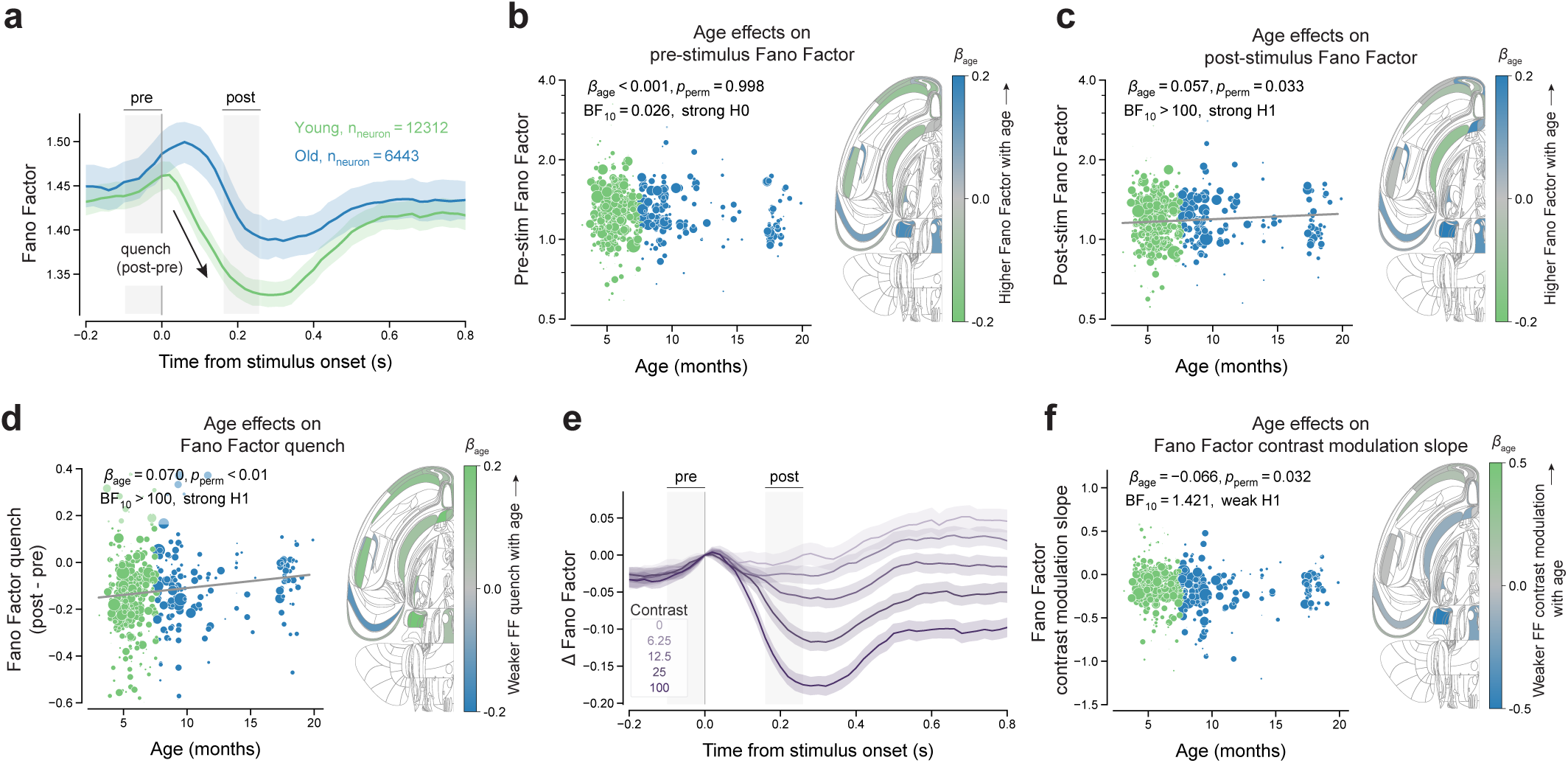
Age effects on global mean-subtracted Fano factor and contrast modulation. (a) Time courses of the overall mean-subtracted Fano factor, aligned to stimulus onset. Thick lines show the mean within each age group; shaded areas show 95% confidence intervals obtained by pooling neurons within each age group. The analysis included n = 18,755 neurons from n = 503 insertions and n = 149 mice. Grey areas indicate the pre-stimulus (-100 to 0 ms) and post-stimulus (160 to 260 ms) windows. For left panels in b, c, d and f, dots show insertion-level summaries for visualization, with dot size indicating the number of neurons recorded in each insertion; statistical analyses and fitted regression lines were based on single-neuron metrics. All statistical conclusions were based on Bayes Factors (BF_10_); two-sided permutation p-values are unadjusted for multiple comparisons and are reported as complementary statistics (see Methods). (b) Left: relationship between pre-stimulus Fano factor and mouse age. *p*perm=0.99800. Right: a flatmap representation showing region-specific age effects on pre-stimulus Fano factor. The colour indicates the slope of age. White areas were not covered in our recordings. (c) Same as (b) but showing post-stimulus Fano factor. *p*perm=0.03297. (d) Same as (b) but showing Fano factor quenching (post-stimulus Fano factor minus pre-stimulus Fano factor). *p*perm=0.00799. (e) Contrast modulation of Fano factor quenching. Saturation indicates stimulus contrast level; thick lines show the mean within each stimulus contrast level; shaded areas indicate 95% confidence intervals. (f) Same as (b), but for contrast modulation slope. Slopes were computed for each neuron by fitting a linear relationship between stimulus contrast and the change in baseline-corrected mean-subtracted Fano factor. *p*perm=0.03197.

Older animals showed attenuated variability quenching following stimulus onset. Neural variability, as measured by the Fano factor, typically decreases upon stimulus onset across various cortical regions and species^25–29^, also called variability quenching. We quantified a quench index as the difference between post- and pre-stimulus Fano factors. We found quenching across the brain attenuated with age, becoming less strongly negative (Figure 4d, left panel). Our region-specific analyses revealed age-related attenuation in variability quenching in visual and motor cortex (VISp+pm, MOs), striatum (LS, ACB) and thalamic area LP (Figure 4d, right panel; Supplementary Figure 15).

Contrast-modulated Fano factor quenching changed weakly with age. Similar to the contrast modulation of firing rate, we examined how the Fano factor varied with stimulus contrast (Figure 4e, Supplementary Figure 16). We again computed a contrast-modulation slope for each neuron by fitting a linear relationship between stimulus contrast levels and the change in baseline-corrected Fano factor (or quench index), revealing a weak, negative effect of age on contrast modulation overall (Figure 4f, left panel). Region-specific analyses identified age-related decreases in the contrast-modulation slope in midbrain regions (SCm, MBm), and strong evidence for no age-related changes in the striatum (LS) and thalamus (LP) (Figure 4f, right panel; Supplementary Figure 17b).

Lastly, although we here focus on animals’ chronological age, neural differences may also be driven by other factors correlated with age. Here, we specifically investigate two sources of interest: training duration and movement.

### Age effects on neural variability persist after accounting for training duration

Age-related effects measured at recording could, in principle, be entangled with differences in training history, as mice vary substantially in learning speed. Although individual training duration explains only a small fraction of variance in age at recording within our dataset (Supplementary Figure 2), and we did not observe slower learning in older mice (Supplementary Figure 1), training duration in the IBL task varies systematically across animals and laboratories^64^. Thus, the neural differences we observe could reflect older animals having followed a longer training trajectory rather than chronological age per se. To address this alternative explanation, we quantified each animal’s training duration and tested its association with our neural metrics.

To directly compare the two predictors, we fitted extended models including both age and training duration for each metric. Across metrics, the evidence for age effects was essentially unchanged after adding training duration (Supplementary Table 1 and 2): pre- and post-stimulus firing rates, post-stimulus Fano factor, and FF quenching all retained strong evidence for an age effect in both the age-only and extended models.

Training duration showed limited, metric-specific contributions and did not change conclusions about effects of age (Supplementary Table 2). Training duration provided strong additional evidence for post-stimulus firing rate and moderate additional evidence for pre-, post-stimulus Fano factor, but did not reduce the evidence for age. For several metrics (e.g., pre-stimulus firing rate, firing rate contrast-modulation slope, FF contrast-modulation slope), the extended models instead provided strong evidence against a training-duration effect. Critically, for Fano factor quenching, the joint models continued to favour an age effect while providing moderate evidence for the absence of a training-duration effect, indicating that reduced neural variability quenching in older animals is not explained by differences in training duration.

### Age effects are robust to video-based movement covariates

Neural activity is modulated by movement^65–67^, including in well-trained animals^68^, motivating a control analysis to test whether the reported age-related neural differences could be explained by movement. Using IBL right-camera videos, we extracted time-resolved paw speed (task-relevant) and nose-tip speed (less task-specific) (Supplementary Figure 18 a,d) from DeepLabCut tracking. Across QC-filtered sessions with valid right-camera tracking (n = 343), neither movement measure showed a systematic association with age in the pre- and post-stimulus windows used to define our neural metrics (Supplementary Figure 18 b-c, e-f). We next incorporated both movement measures (paw speed and nose-tip speed) as covariates in extended models for each neural metric (Supplementary Table 3). Movement covariates provided additional explanatory power in several models, but the qualitative evidence for age effects was broadly preserved after including them. Only two metrics with initially weak-to-moderate evidence were sensitive to the inclusion of movement covariates (firing rate contrast modulation: moderate H0 to weak H1; Fano factor contrast modulation: weak H1 to moderate H1). All other metrics retained the same qualitative conclusion.

To further test whether the age-related neural effects could be accounted for by movement variability beyond average movement speed, we performed an additional control analysis using the coefficient of variation (CV) in trial-to-trial right-paw and nose-tip speed within the same pre- and post-stimulus windows. Session-level movement variability did not show consistent evidence for a systematic association with age, across measures and time windows (Supplementary Figure 19). Although these covariates explained additional variance in several cases, adding these movement-variability measures as covariates to the extended models left the main age-related pattern broadly unchanged (Supplementary Table 4).

Together, these analyses suggest that while video-derived movement explains additional variance in firing-rate and variability metrics, the reported age-related neural effects persist after accounting for both movement magnitude and movement variability.

## Discussion

In this study, we investigated age-related changes in behaviour and neural activity using large-scale extracellular recordings in mice. Older mice performed slightly worse (showing increases in absolute bias, perceptual thresholds, and lapse rate) and showed increased variability in their response times. We observed global changes in single-neuron baseline firing rates, post-stimulus Fano factors, and an age-related reduction in the amplitude of stimulus-induced variability quenching. Older animals showed higher overall firing rates across areas of visual and motor cortex, striatum, midbrain, and hippocampus, but lower firing rates in thalamic areas. Age-related attenuation in stimulus-induced variability quenching was most prominent in visual and motor cortex, striatum, and thalamic area LP.

Older animals’ slightly worse task performance could be due to a combination of peripheral (sensory, motor) and cognitive sources. Age-related decline in sensory processing, particularly in vision, is well-documented across species^69–73^. However, visual deficits in C57BL/6 mouse strains start manifesting most dramatically after 20 months, beyond the age range of this study^74^. Crucially, a sole failure to process the sensory stimulus (due to poor eyesight or the inability to hear the stimulus onset tone) would lead to an overall slowing of responses in older animals, which we did not observe. Our findings thus suggest that cognitive and decision factors also contribute to the behavioural and neural effects we observed.

The more variable RTs in older mice align with a well-established pattern in human ageing research: in human cohorts of a roughly equivalent age range, response time variability increases in a large range of cognitive tasks^5,49,51^. The same age-related increase in RT variability in mice provides evidence for cross-species consistency in behavioural variability, highlighting a potentially fundamental aspect of behavioural ageing conserved across mammalian species.

The slight reduction in overall neural yield observed in older mice may reflect a combination of technical (recording-related) and biological factors. From a technical perspective, increased tissue and dural rigidity during probe insertion may increase superficial damage and reduce the number of well-isolated units; however, the regional specificity of the effect suggests that insertion-related factors alone are unlikely to account for the observed pattern. Biologically, ageing has been associated with modest and region-specific neuronal loss in some cortical and subcortical regions^75–77^. More broadly, ageing is accompanied by cytoarchitectural and structural remodeling, including changes in cellular organization, dendritic morphology, and synaptic integrity^78,79^. Such age-related changes may reduce the number of stable, well-isolated units obtained in large-scale extracellular recordings, even in the absence of pronounced neuron loss.

Our investigation across 16 brain regions showed age-related increases in single-neuron firing rates. This finding is in line with several studies reporting increased firing rates in aged primary sensory cortices, including cat V1^19^ and macaque V1^20^, and in specific cell types in monkey prefrontal cortex, including pyramidal cells^18^. We also found that post-stimulus Fano factors increased with age, consistent with previous findings from anesthetized rhesus monkeys^21^. Future work could also examine complementary measures of neural variability beyond Fano factor, such as spike train irregularity^80,81^, to test the robustness and specificity of these effects. With our broad sampling of neurons across cortical and subcortical brain regions, surpassing that of prior studies, our findings indicate that age-related changes in firing rates and neural variability may be present across species, brain regions, and behavioural states.

We also observed significant age-related differences in stimulus-induced variability quenching. Previous studies found that neural variability decreases (quenches) upon stimulus onset in various cortical regions and species, including a wide range of cortical regions of monkeys^25,29^, the PPC of rats^27^, and the olfactory cortex of both mice^26^ and rats^28^, although thalamic nuclei did not show stimulus-induced quenching of neural variability^29^. Here, we found an age-related reduction in variability quenching, most prominently in the visual and motor cortex (VISp+pm, MOs), striatum (LS, ACB) and thalamic area LP. In older animals, attenuated (i.e., less strongly negative) variability quenching may reflect a lower temporal precision of task-relevant neural processes underlying decision-making that in turn drives greater trial-to-trial variability in response times. Future work could further explore and quantify how these single-neuron changes in variability propagate to neural circuits and networks, and lead to the computations that drive age-related changes in behaviour.

Changes in single-neuron variability may arise from various biological mechanisms. Age-related degradation of inhibitory and neuromodulatory systems^5,51,82,83^ may cause reduced gain modulation, shorter intrinsic neural timescales^84,85^, and lower temporal precision of single-neuron responses. These could in turn be reflected in pairwise noise correlations^23,86^, neural population dynamics^16,87,88^, and cortical state (e.g., as reflected in multi-unit activity power spectra)^62,89^. Future work on such population-level measures may further reveal how age-related neural variability across scales impacts behaviour.

While we here used animals’ chronological age at the time of their neural recording, this does not imply that age itself is the best predictor of the neural effects we observed. More broadly, the correlational nature of these analyses highlights a general challenge in interpreting ageing effects: chronological age covaries with multiple behavioural and biological factors that may jointly shape neural activity and variability, and these influences cannot be fully disentangled in the current dataset. Specifically, other factors such as training duration, movement patterns during task performance, or different decision strategies may give rise to different task strategies or different computational mechanisms that produce similar overt task behaviour. While we controlled for video-derived movement covariates (paw, nose-tip), future work could further refine such integrated predictors using richer behavioural characterizations^90^, for example motion-energy–based video features or higher-dimensional, data-driven representations of behavioural modules^91,92^. Accordingly, a range of biological, behavioural, and life history factors could be further integrated to compute an individual animal’s brain age^93^ as opposed to their chronological age, which may reveal higher predictive power in explaining between-animal differences in behavioural and neural variability.

In conclusion, we present a survey of age-related neural variability in >18,000 neurons across 16 brain regions in behaving mice, complementing the previously released brain-wide map recording in younger animals^45^. Our findings replicate earlier work on neural correlates of ageing in other species and brain areas, and show how multi-area neural recordings in behaving animals can help uncover regional specificity of age-related changes in neural variability. Lastly, we provide a well-curated, open dataset in a standardized data format^64^ that further allows investigating the computational mechanisms through which neural variability gives rise to age-related cognitive decline.

## Methods

### Data

This study included a previously released dataset (the brain-wide map dataset)^45^ from the International Brain Laboratory (IBL) and our new release (the lifespan dataset). Both datasets contain extracellular recordings of mice performing a visual decision-making task, collected using the same standardized protocols^94,95^. All experimental procedures for the newly collected lifespan dataset were conducted in accordance with local laws and approved by the institutional IACUC at Cold Spring Harbor Laboratory (licences 1411117 and 19.5).

### Animals

We included 130 mice from a previously released dataset (C57BL/6, 89 male, mean age = 6.64 months, range 3.10–15.13 months) and 19 mice from a newly recorded dataset (C57BL/6, 11 male, mean age = 16.58 months, range 10.58–19.90 months). All mice were housed and cared for following standardized protocols^43^. Age was calculated on the day of electrophysiological recording. Across mice, age at recording varied due to differences in (1) age at the start of behavioural training, (2) training duration, and (3) non-training intervals between training start and the first recording (e.g., experimental or logistical constraints). A decomposition of these sources indicates that variability is dominated by differences in age at training start (Supplementary Figure 2). After quality control, 149 mice were included in the analysis. For visualization, mice were categorized based on their age at recording into a young group (N = 97, mean age 5.60 months) and an old group (N = 52, mean age 12.07 months), using 7.6 months (mean age in the dataset) as the age cutoff (Figure 1b). Age was treated as a continuous variable in all statistical analyses.

### Behavioral task

Mice were trained to perform a standardized visual decision-making task^43^. In this task, mice had to decide the location of a visual stimulus presented on the screen in front of them. Specifically, each trial began when the mouse held the wheel still for 0.4-0.7 seconds. Then, an auditory cue (a 100-ms tone, 5 kHz sine wave) and a visual stimulus (Gabor patch) were presented on either the left or right side of the screen. Mice indicated the stimulus location by turning the response wheel to bring the stimulus to the centre of the screen. They had up to 60 seconds to make a response. The contrast level of the visual stimulus varied across trials. There were five different contrast levels (%): 100, 25, 12.5, 6.25, 0. Combining the five contrast levels with the two stimulus sides (left, right) resulted in nine conditions: -100, -25, -12.5, -6.25, 0, 6.25, 12.5, 25, 100 (negative for left stimuli, positive for right stimuli). Correct responses were rewarded with sugar water, while incorrect responses were followed by a noise burst and a longer inter-trial interval. Stimulus probabilities were fixed within each block: sessions began with an unbiased block (50:50 left vs. right) and then alternated between blocks biased toward the left (80:20) and toward the right (20:80). The task was based on the standardized IBL visual decision-making task introduced in the original task paper^43^. Animals were trained according to the IBL training protocol^96^. We here provide details relevant to the present study.

### Behavioural measures

#### Psychometric curve

Psychometric curves illustrate how the probability of a rightward choice changes with stimulus position and stimulus contrast. For each session, responses were fitted with a parametric error function using a maximum likelihood procedure:

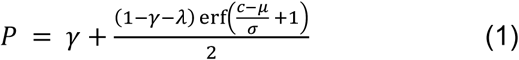

*P* is the probability of rightward choice;

*c* is the contrast level;

*γ*, *λ*, *σ*, *μ* are fitted parameters:

*γ*: lapse low (the lapse rate for left stimuli);

*λ*: lapse high (the lapse rate for right stimuli);

*σ*: the decision threshold;

*μ*: the response bias (horizontal shift of the curve);

For details on the fitting process and parameter computation, see the IBL protocol^96^. The fitted psychometric parameters were obtained for each session. In the statistical analyses, we examined ageing effects by correlating the absolute bias |*μ*|, threshold *σ*, and mean lapse 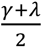 to the age of mice.

#### Chronometric curve and RT variability

Chronometric curves show how response time varies with stimulus position and contrast. RT was defined as the time between stimulus onset (stimOn_times, when the visual stimulus appeared on the screen) and response (response_times, recorded either after 60 seconds, i.e., timeout, or when the rotary encoder indicated the stimulus reached ±35° azimuth). A chronometric curve was computed for each session.

To assess RT variability, we computed the coefficient of variation (CV) for each session.

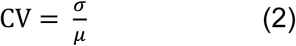

Where:

*σ* = standard deviation of response times

µ = mean response time

### Neuropixels recordings

Extracellular recordings were acquired using Neuropixels probes^47^. For a detailed description of the animal surgery, apparatus, and recording procedure, see Appendix 2 and 3^44^. Up to two probes were inserted during each recording session.

### Spike sorting

All sessions were spike sorted using ibl-sorter (version 2.35.0)^97^, matching the version used for the original IBL dataset processing^45^. The ibl-sorter code is publicly available^98^.

### Quality Control

Before analyzing the data, we applied a set of inclusion criteria (at the session, insertion, and neuron level) to ensure data quality. These criteria were primarily based on previously established IBL criteria^45^, with additional requirements for single-neuron quality. These additional requirements were included to ensure that Fano factor estimates were computed from neurons with stable activity, as this metric is particularly sensitive to unstable firing. The inclusion and exclusion criteria are described below.

#### Sessions and insertions

(1) Sessions with more than 400 trials and at least 90% accuracy on 100% contrast trials were included. (2) There must be at least 3 trials with incorrect responses (after applying the trials filter below). (3) Sessions had to pass the hardware tests, as defined at https://int-brain-lab.github.io/iblenv/_autosummary/ibllib.qc.task_metrics.html. (4) Sessions were visually inspected to exclude those which have pronounced instability of recordings (drift), epileptiform activity, noisy channels, and artifacts (following the Recording Inclusion metrics and Guidelines for Optimal Reproducibility (RIGOR) criteria^44^). (5) Insertions must have resolved alignments (see Appendix 6^44^ for definition). (6) Insertions were also required to have spike-sorted data processed using the same version of the ibl-sorter algorithm (version 2.35.0).

#### Trials

To control for differences in trial counts across sessions, we included only the first 400 trials of each session. Additional exclusion criteria were applied: (1) trials missing any of the following events—choice (response type), probabilityLeft (stimulus side probability), feedbackType (positive for correct, rewarded responses; negative for incorrect or timed-out trials), feedback_times (time of feedback), stimOn_times (stimulus onset), or firstMovement_times (the time of the first detected movement with sufficient amplitude); and (2) trials with RTs outside the range of 0.08–2 s^45^.

#### Neurons

Neurons included in the analyses met the following criteria: (1) passed three single-unit quality control (QC) metrics—a refractory period violation metric, a noise cutoff metric, and a median amplitude threshold—based on the computed single-unit metrics of RIGOR^44^; (2) had an average firing rate greater than 1 spike/s; and (3) presence ratio exceeded 0.95. Final analyses were further restricted to 16 ROIs (see Table 1).

ROIs include a combination of areas from the Beryl parcellation^45^. We chose this set of insertion targets for a combination of scientific (earlier age-related degradation in more frontal regions), statistical (inclusion of the repeated site that was used to confirm reproducibility across labs^44^), and practical reasons (surgical accessibility).

Table 2 shows the number of sessions and insertions that survived after each quality control criterion was applied. After quality control, we included 367 recording sessions and 503 insertions in the final analyses.

### Fano factor

Trial-by-trial neural variability was quantified using the Fano factor, defined as the variance of spike counts across trials divided by the mean spike count:

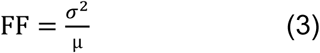

Where:

*σ*^2^ = variance of spike count across trials

µ = mean spike count across trials

Because neural responses can differ across stimulus conditions, we first divided trials into nine stimulus conditions (side × contrast: −100, −25, −12.5, −6.25, 0, 6.25, 12.5, 25, 100%) to prevent differences in mean responses from artificially inflating variance estimates.

#### Mean-subtracted Fano factor

To ensure that our Fano factor analyses were not biased by differences in mean firing rates across conditions, we adapted the mean-subtraction method^63^ to pool trials while controlling for condition-specific differences in mean firing rates. Specifically, for each trial and time window, we subtracted the condition-specific mean spike count to obtain a residual. Mean-subtracted Fano factors were then computed from the variance of these residuals using a sliding-window approach (width = 100 ms, step size = 20 ms) for each neuron.

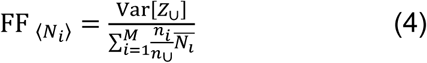

It is defined as the variance of the union of residuals from all conditions *Z*_∪_, divided by the weighted average of the mean spike count for the conditions. where *n*_∪_ is the total number of trials across all *M* conditions, *n_ii_* and 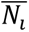 are the number of trials and the mean count for the *i* -th condition, respectively. Statistical tests were performed on these mean-subtracted Fano factors.

### Statistical tests

We assessed the effect of age on each behavioural and neural metric. Behavioural metrics (absolute bias, threshold, mean lapse rate, median RT, and RT variability) were computed at the session level. Neural metrics included neural yield, firing rate, contrast modulation of firing rate, Fano factor, Fano factor quench, and contrast modulation of Fano factor. For each neuron, contrast-modulation slopes were obtained by fitting a linear regression to the change in firing rate (Δ firing rate, post-minus pre-stimulus window) or Fano factor (Δ Fano factor, post-minus pre-stimulus window) across all contrast levels. Neural yield was defined and analysed at the probe-insertion level, whereas all other neural metrics were computed at the single-neuron level, and all statistical analyses were performed on these neuron-level metrics.

To test for age-related effects, we used the age-related slope, *β*age, from a general linear model (Gaussian family, identity link, using a generalized linear modelling framework) as the test statistic. Strictly positive and right-skewed measures (median RT, psychometric threshold, firing rates, Fano factors) were analysed on a log scale to reduce skewness and stabilise variance. In contrast, RT variability, signed change indices (e.g. FF quench and contrast modulation metrics), nonnegative metrics with substantial mass near zero (absolute bias), and bounded proportions (mean lapse) were analysed on their natural scale, since log transforms are undefined at zero and can introduce artificial heavy lower tails when values are concentrated near zero. We inspected residual Q–Q plots for representative models; residuals for log-transformed metrics showed approximate normality with mild tail deviations, whereas bounded and difference metrics exhibited somewhat heavier but unimodal tails. Importantly, we verified that our conclusions did not change depending on the log-scaling of these variables.

The following metric-specific transformations and covariates were used

**Behavioural metrics:** median RT and psychometric threshold were analysed as log(*y*), with age as the only predictor. RT variability, absolute bias, and mean lapse were analysed on their natural scale, with age as the only predictor.

**Neural metrics:** neural yield was analysed on its natural scale with age and brain region as predictors. Pre- and post-stimulus firing rates were analysed as log(*y*), with age, brain region, absolute signed contrast, and trial count as predictors. Firing rate contrast modulation slope was analysed on its natural scale with age and brain region as predictors. Pre- and post-stimulus Fano factors were analysed as log(*y*), with age, brain region, and trial count as predictors. Fano factor quench was analysed on its natural scale with age, brain region, and trial count as predictors. Fano factor contrast modulation slope was analysed on its natural scale with age and brain region as predictors.

#### Bayes Factors

We computed Bayes Factors (BF_10_), using the BayesFactor^99^ and Pingouin^100^ packages, to quantify evidence for including mouse age as a predictor in each model. BF_10_compares models with and without age, with values >1 indicating support for an age effect (i.e., age modulates the behavioural or neural metric) and <1 favouring the null. Statistical conclusions throughout the manuscript were based on BF_10_.

For visualization purposes, regression lines are plotted only when the BF_10_ provides strong (BF_10_ > 10) or moderate (10 ≥ BF_10_ > 3) evidence in favour of the alternative hypothesis (H1). While the choice of these thresholds is inherently arbitrary, we adopted them to ensure that plotted regression lines correspond to effects with at least moderate statistical evidence.

#### Permutation tests

We additionally computed permutation-based *p*-values which are shown alongside the Bayes Factors. Note that all conclusions are based on the Bayes Factors, which provide a primary graded measure of evidence for or against an effect. Permutation tests additionally account for the hierarchical data structure by shuffling age labels at the session level. In omnibus analyses, both methods led to the same conclusions. For region-specific results, we highlighted regions with strong or moderate evidence for the alternative (H1), as well as those with strong evidence for the null (H0).

To compute permutation-based *p*-values, the null hypothesis was that there is no association between the metric and mouse age. To generate a null distribution for each analysis, we performed a permutation procedure (n = 10,000 iterations for behavioural metrics; n = 2,000 iterations for neural metrics). Importantly, for the neural metrics (except for neural yield), we randomly shuffled the mouse age labels across recording sessions in each iteration while maintaining the grouping of neurons recorded within each session. This session-based shuffling preserved the inherent dependencies among neurons recorded simultaneously. After each shuffle, we recalculated the test statistics for the specific metric being examined. The *p*-value for each metric was then calculated as the proportion of the permuted age slopes whose absolute values were equal to or greater than the absolute value of the observed slope from the original data. This two-tailed *p*-value represents the probability of observing a relationship as strong as, or stronger than, the one found, under the null hypothesis of no relationship between the metric and age.

#### Extended models including age and training duration

To assess whether our age effects could be explained by training duration, we fitted extended models for key metrics that included both age and training duration as predictors simultaneously. Training duration was defined as the number of training days until the mouse first passed the standardised behavioural criterion and was expressed in years to match the age scale used in the model. All other modelling choices (predictors, covariates, and log-transform conventions) were identical to those described above.

In each extended model, we quantified evidence for including each predictor (age, training duration) using Bayes Factors that compared otherwise identical models with and without the respective term. This allowed us to test whether age effects persisted after accounting for training duration, and whether training duration explained additional variance beyond age.

#### Extended models including age and movement

We used IBL right-camera videos to derive movement covariates from DeepLabCut tracking of two representative keypoints: right paw (task-relevant) and nose tip (task-irrelevant) (Supplementary Figure 18 a,d). From the tracked x/y coordinates, we computed frame-to-frame speed (coordinates normalized by camera resolution and scaled by the camera sampling rate), and aligned the resulting speed time courses to stimulus onset. We summarized movement by computing median speed within the same pre- and post-stimulus windows used for neural metrics (pre: −100 to 0 ms; post: 160 to 260 ms), yielding pre- and post-stimulus movement measures for both paw and nose tip. To quantify movement variability, we additionally computed the coefficient of variation (CV; standard deviation / mean) in trial-to-trial paw and nose-tip speed within the same pre- and post-stimulus windows.

Sessions were required to pass IBL extended quality control for the right camera (videoRight and dlcRight). This yielded n = 343 sessions (out of total 367 sessions) with valid right-camera movement measures. We then fitted extended statistical models for each neural metric by adding either movement-speed covariates or movement-variability covariates to the age-only models. Movement covariates were matched to each neural metric by time window and contrast structure. Metrics defined in a single window used the corresponding pre- or post-stimulus movement measure, whereas metrics spanning both windows included both pre- and post-stimulus covariates. For neural metrics retaining contrast-specific values (pre- and post-stimulus firing rate), movement measures were computed separately for each session and contrast condition; for neural metrics not retaining contrast (contrast modulation of firing rate, pre- and post-stimulus Fano factor, Fano factor quench, and contrast modulation of Fano factor), movement measures were then aggregated across contrasts to yield one value per session, window, and feature. We applied the same matching scheme in the movement-variability control models. All other modelling choices (predictors, covariates, log-transform conventions, and inference procedures) were identical to those described above.

## Data availability

The data analysed in this study are publicly available through the International Brain Laboratory website (https://www.internationalbrainlab.com/data), under the tag ‘2025_Q3_Zang_et_al_Aging’. Instructions for accessing these data are provided in the GitHub repository associated with this study (https://github.com/Fenying-Zang/Ageing_behavioral_and_neural_variability). The analysis results underlying the figures are also available in the same repository.

## Code availability

Data analyses were performed using custom Python scripts in a reproducible Python environment. All code required to reproduce the figures and analyses is available at https://github.com/Fenying-Zang/Ageing_behavioral_and_neural_variability, and has been archived on Zenodo^101^.

## Acknowledgements

AEU thanks Graham Wildt and John Pisciotta for assistance with laboratory safety procedures, Rachel Rubino for excellent animal care advice, and Joao Couto for help with craniotomy surgeries. Nathaniel Miska contributed recordings from two aged animals. Henk van Steenbergen, Bryant Jongkees, Steven Miletić, and Jacqueline Zadelaar provided valuable feedback on statistical analyses. Sander Nieuwenhuis offered helpful comments on the first draft. The IBL development team, especially Olivier Winter, provided crucial technical support with spike sorting and data release. Pranav Rai provided helpful code review. Peter Dayan and Kenneth Harris provided insightful suggestions throughout the project, and we thank the CoCoSys lab for discussions and feedback.

## Author contributions

F.Z., A.K.C. and A.E.U. conceptualized the study. A.K. and A.E.U. contributed to the methodology for animal training, and A.E.U. contributed to the methodology for neural recordings. F.Z. and A.E.U. performed the formal analysis. F.Z., A.K., S.F. and A.E.U. contributed to data curation. The IBL provided software. F.Z. and A.E.U. contributed to visualization. F.Z. and A.E.U. wrote the original draft. F.Z., A.K., S.F., A.K.C. and A.E.U. reviewed and edited the manuscript. A.K.C. and A.E.U. supervised the study. F.Z., IBL, A.K.C. and A.E.U. acquired funding. Please see the Contribution Table (Supplementary Figure 20) for additional details.

