## Supplemental materials for "Age-related changes in behavioural and neural variability in a decision-making task"

### Supplementary Figures

**General plotting and statistical conventions for Supplementary Figures.** Unless otherwise stated, age effects were estimated using general linear models with age as a continuous predictor. All statistical conclusions were based on Bayes Factors ($\mathrm{BF}_{10}$); two-sided permutation p-values are unadjusted for multiple comparisons and are reported as complementary statistics (see Methods). Exact statistical values are shown in the figures or provided in the Source Data file. Regression lines are plotted only when the Bayes Factor indicates strong or moderate evidence for H1. Shaded areas and error bars indicate 95% confidence intervals.

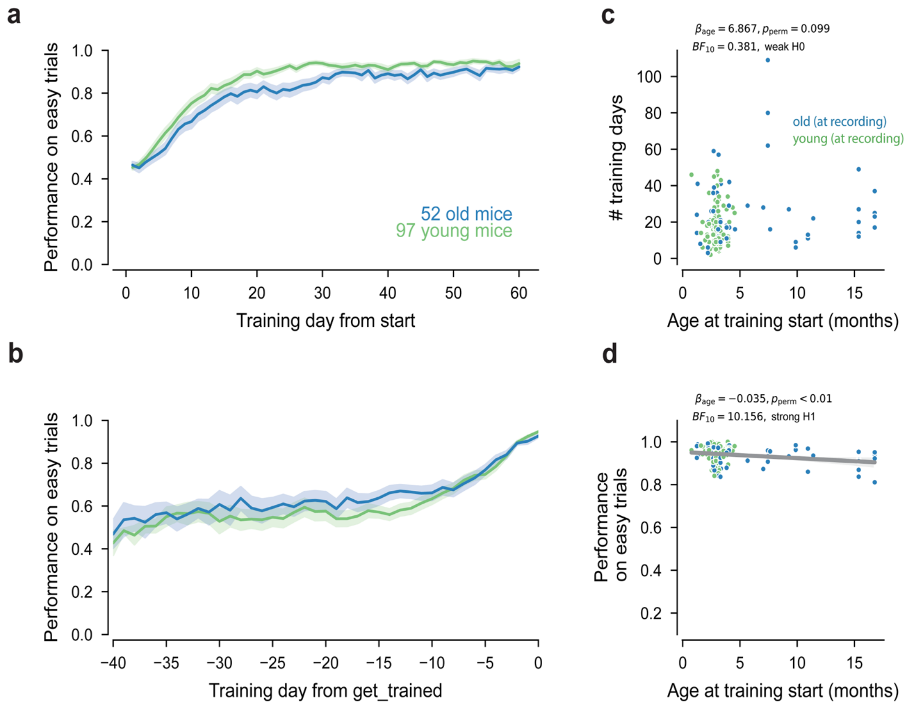


**Supplementary Figure 1, related to Figure 1. Training progression in the behavioral task.** All panels include the same mice unless otherwise stated: n = 149 mice (52 old mice and 97 young mice, age group defined at recording). **(a)** Learning curves from the start of training. Age group split was based on the age at recording, following the same convention as in other main figures. Solid lines show mean performance across mice; shaded areas indicate 95% confidence intervals. **(b)** Training curves aligned to the day animals reached the behavioral criterion ('get_trained', see the training protocol for definition[1]). Solid lines show mean performance across mice; shaded areas indicate 95% confidence intervals. **(c)** Number of training days each mouse took to reach the behavioral criterion. Each point is one mouse. Conclusions are the same when we use a slightly different definition of training duration: number of days, sessions and trials required to reach the behavioural criterion or cumulative days, sessions and trials completed before the first neural recording. **(d)** Performance on easy trials on the day each mouse reached criterion. Each point is one mouse. The training criterion was identical for all mice. Observed differences reflect performance above this threshold within criterion-satisfying sessions on that day.

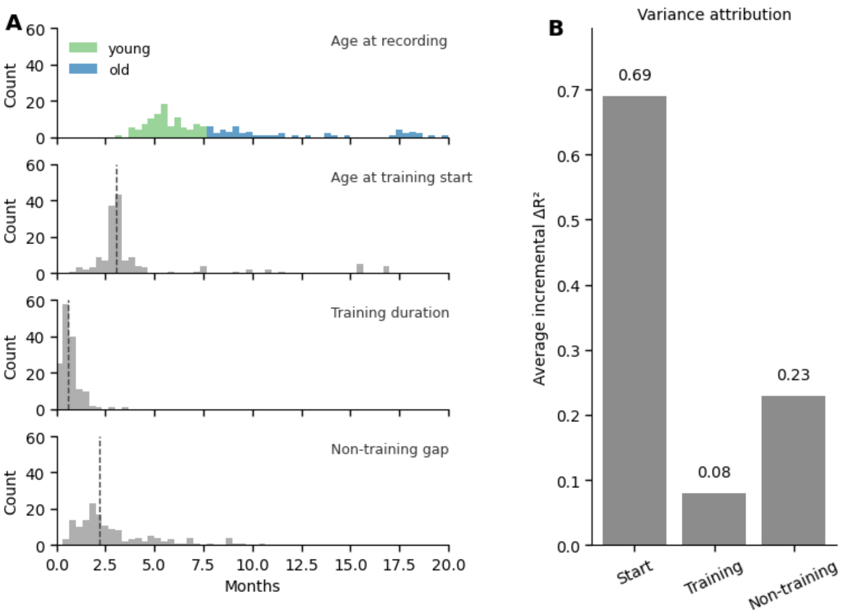


**Supplementary Figure 2, related to Figure 1**. **Components contributing to variability in age at recording.** All analyses used n = 149 mice. **(a)** Distributions (in months) of age at recording and the three components used to decompose age at recording: age at training start, training duration (number of training days), and the non-training gap between training start and the recording. Dashed lines indicate the median. **(b)** Order-invariant variance attribution using average incremental $\Delta R^2$ across all predictor orderings in an OLS model.

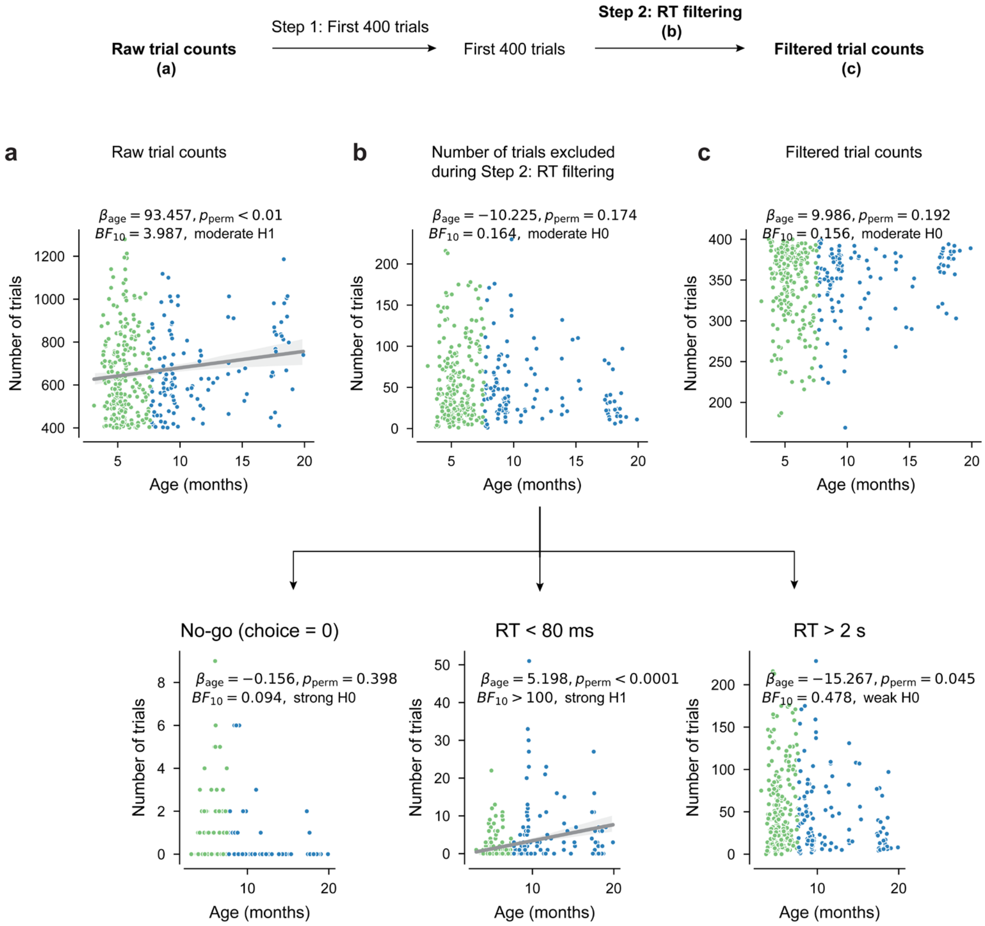

**Supplementary Figure 3, related to Figure 1. Older mice performed slightly more trials in the standardized task.** All panels use session-level data: n = 367 sessions from 149 mice. Each dot represents one session. **(a)** Number of trials as a function of age. **(b)** Number of trials excluded by RT filtering (no-go trials, RT < 80ms trials, and RT > 2s trials). **(c)** Number of trials included as a function of mouse age.

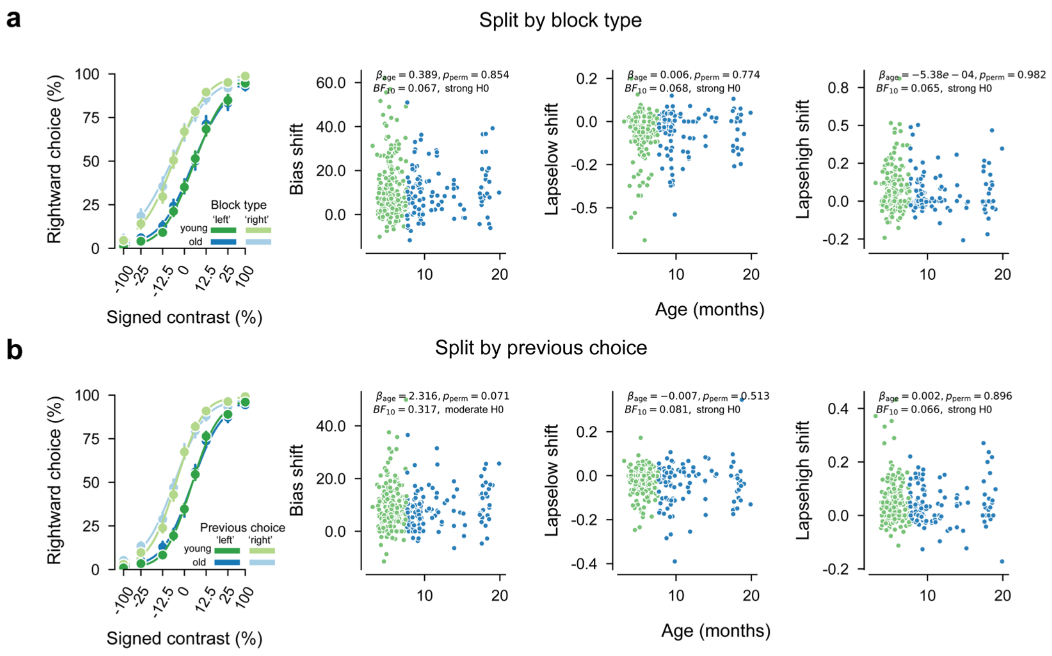


**Supplementary Figure 4, related to Figure 1. No age-related differences in block- or history-dependent choice bias.** All panels use session-level psychometric fits: n = 367 sessions from 149 mice. For psychometric curves, error bars indicate 95% confidence intervals. Each dot in the scatter plots represents one session. **(a)** Trials were split by block type (see Methods), and shifts in psychometric parameters (bias and lapse rate) were quantified between the two block conditions. Left, psychometric curves show mean rightward choice probability as a function of signed contrast, split by age group and block type. No age-related effects were observed for bias shift, low-lapse shift, or high-lapse shift. **(b)** Trials were split based on the previous choice (clockwise and counterclockwise), and shifts in psychometric parameters (bias and lapse rate) were quantified between the two types of previous choices. Left, psychometric curves show mean rightward choice probability as a function of signed contrast, split by age group and previous choice. Again, no age-related effects were observed for bias shift, low-lapse shift, or high-lapse shift.

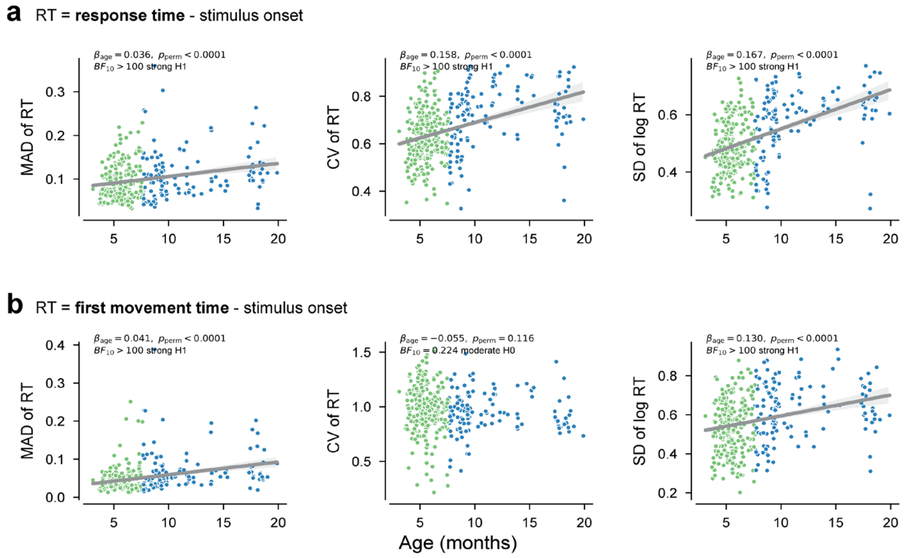


**Supplementary Figure 5, related to Figure 1**. **RT variability measures across two RT definitions and their relationship with age**. All panels use session-level data: n = 367 sessions from 149 mice. Each dot represents one session. Solid lines indicate fitted age effects where plotted; shaded areas indicate 95% confidence intervals. **(a)** RT was defined as the interval between stimulus onset and response time. We examined alternative measures of RT variability that capture different aspects of dispersion. Specifically, we used the median absolute deviation (MAD), a robust metric less sensitive to skew, the coefficient of variation (CV), and the standard deviation (SD) of log-transformed RTs, which stabilizes variance in positively skewed distributions. Scatterplots show the relationship between age and three variability measures: MAD of RT (left), CV of RT (middle), and SD of log-transformed RT (right). **(b)** Same as (a), but for movement initiation time, defined as the interval between stimulus onset and the first movement[2].

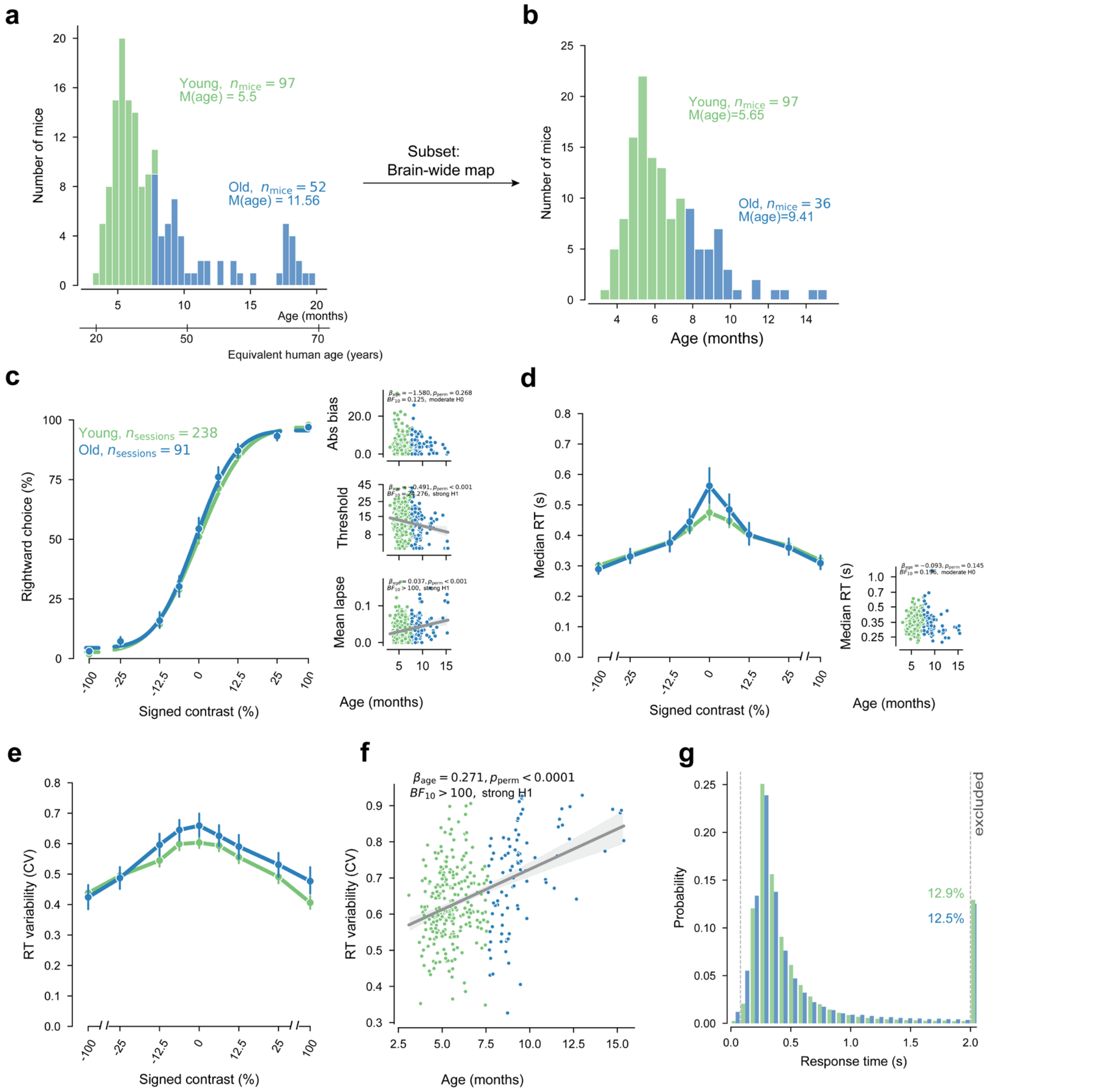


**Supplementary Figure 6, related to Figure 1. Replication of Figure 1 using only mice from the original IBL "brain-wide map" dataset.** Panels, analysis definitions, and plotting/statistical conventions are the same as in main Figure 1, except that analyses were restricted to mice from the original IBL "brain-wide map" dataset[2]: n = 133 mice, 329 sessions.

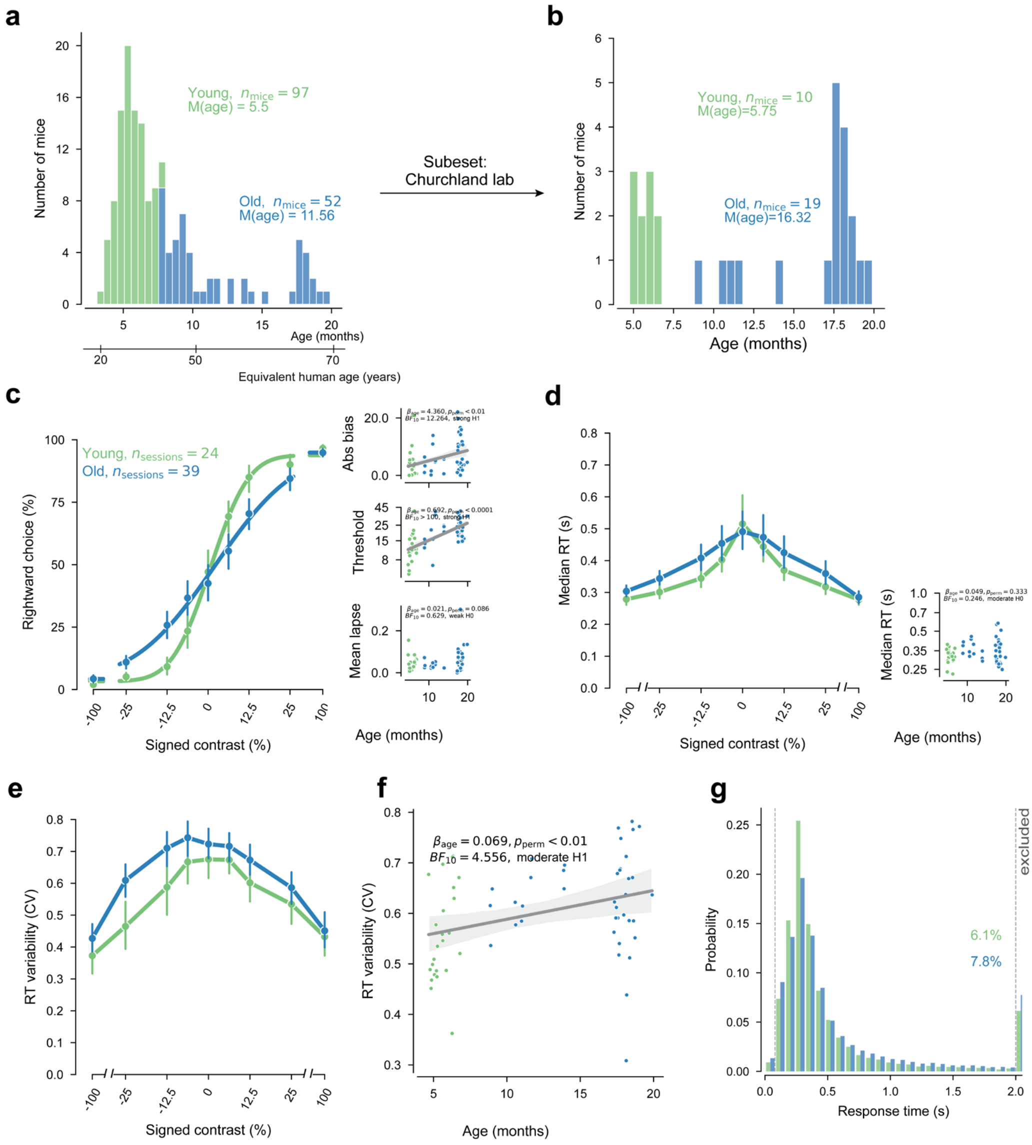


**Supplementary Figure 7, related to Figure 1. Replication of Figure 1 using only sessions recorded in the Churchland lab.** Panels, analysis definitions, and plotting/statistical conventions are the same as in main Figure 1, except that analyses were restricted to sessions recorded in the Churchland lab: n = 29 mice and 63 sessions.

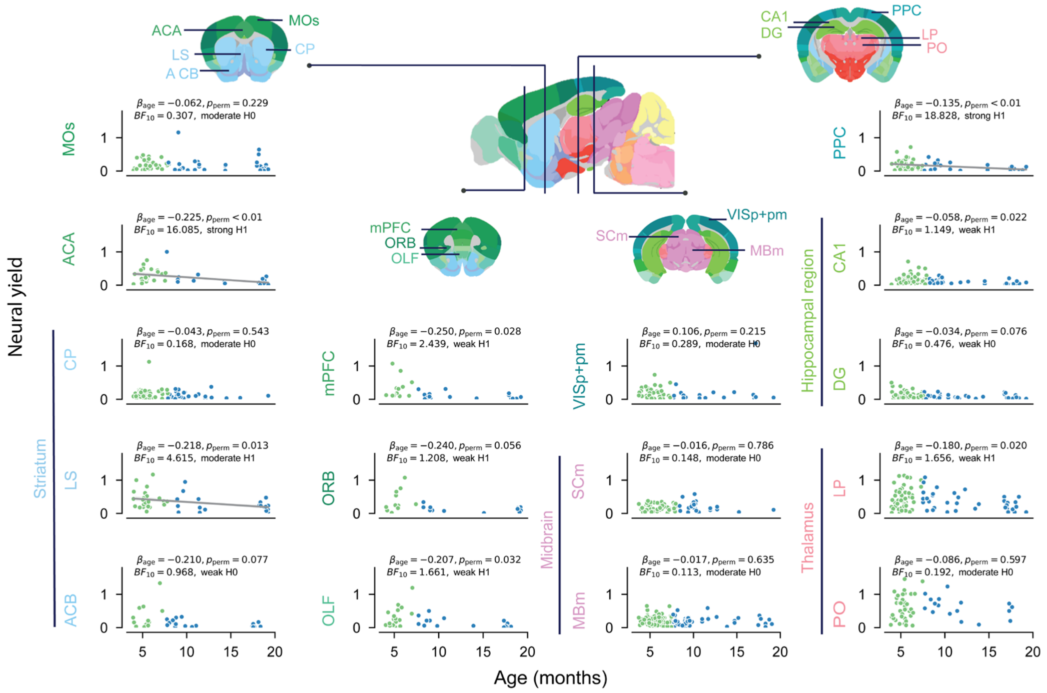


**Supplementary Figure 8, related to Figure 2. Neural yield slightly decreases with age.** Scatterplots show neural yield in each ROI as a function of age. Each dot corresponds to one recording insertion. Statistical analyses for neural yield were performed at the insertion level (yield is inherently defined per insertion). Neural yield is defined as the number of good neurons per good channel (along the length of the Neuropixels probe) within a given region.

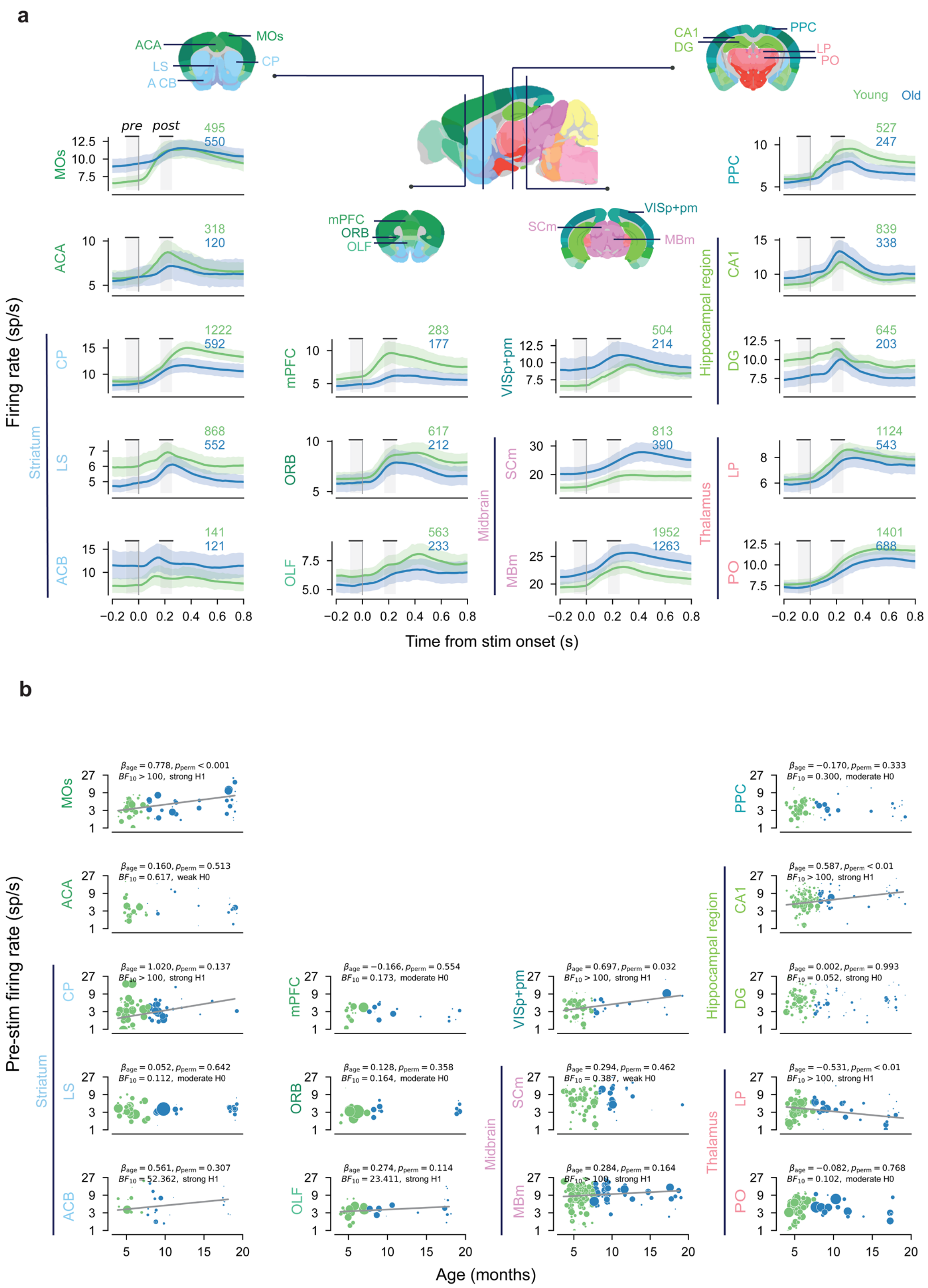


**Supplementary Figure 9, related to Figure 3. Regional specificity of firing rate time courses and pre-stimulus firing rates. (a)** Time courses of mean firing rates in different brain regions, aligned to stimulus onset. Lines show mean firing rates across neurons; shaded areas indicate 95% confidence intervals. ROI-specific n values (number of neurons) are indicated in each panel. **(b)** The relationship between pre-stimulus (−100 to 0 ms relative to stimulus onset) firing rate and mouse age. For visualization, each dot represents the mean across neurons from one insertion, and dot size represents the number of neurons recorded in that insertion. ROI-specific n values are the same as in (a) and indicate the number of neurons included in the statistical analysis.

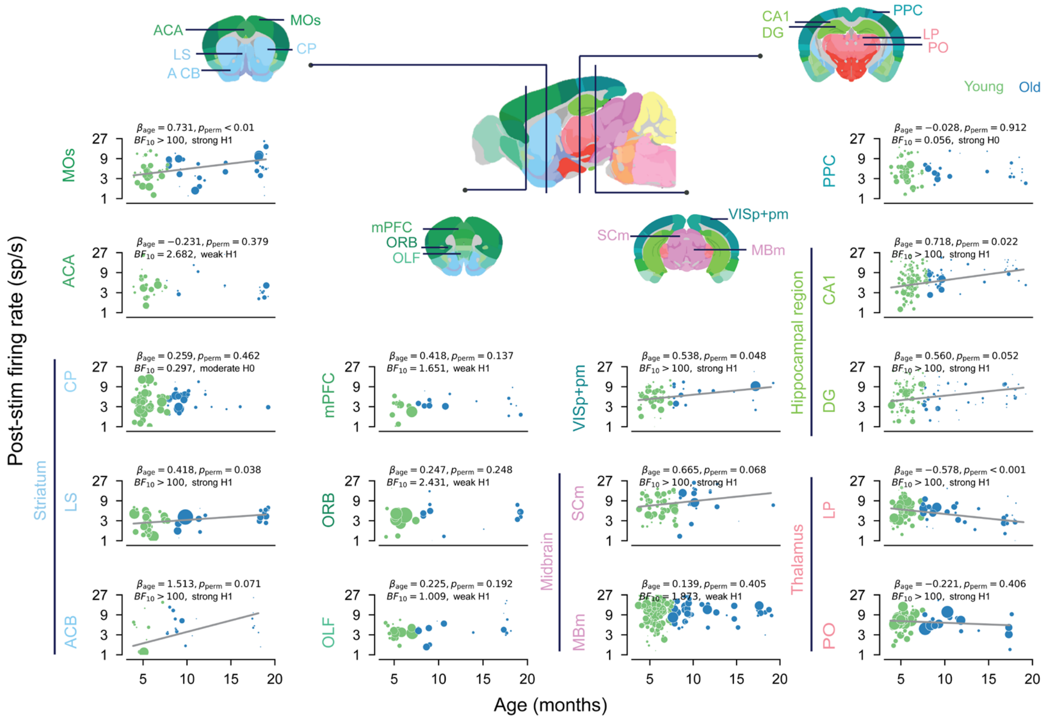


**Supplementary Figure 10, related to Figure 3. Regional specificity of post-stimulus firing rates.** The relationship between post-stimulus firing rate (160 to 260 ms relative to stimulus onset) and mouse age. Visualization, sample-size, plotting, and statistical conventions are the same as in Supplementary Figure 9b.

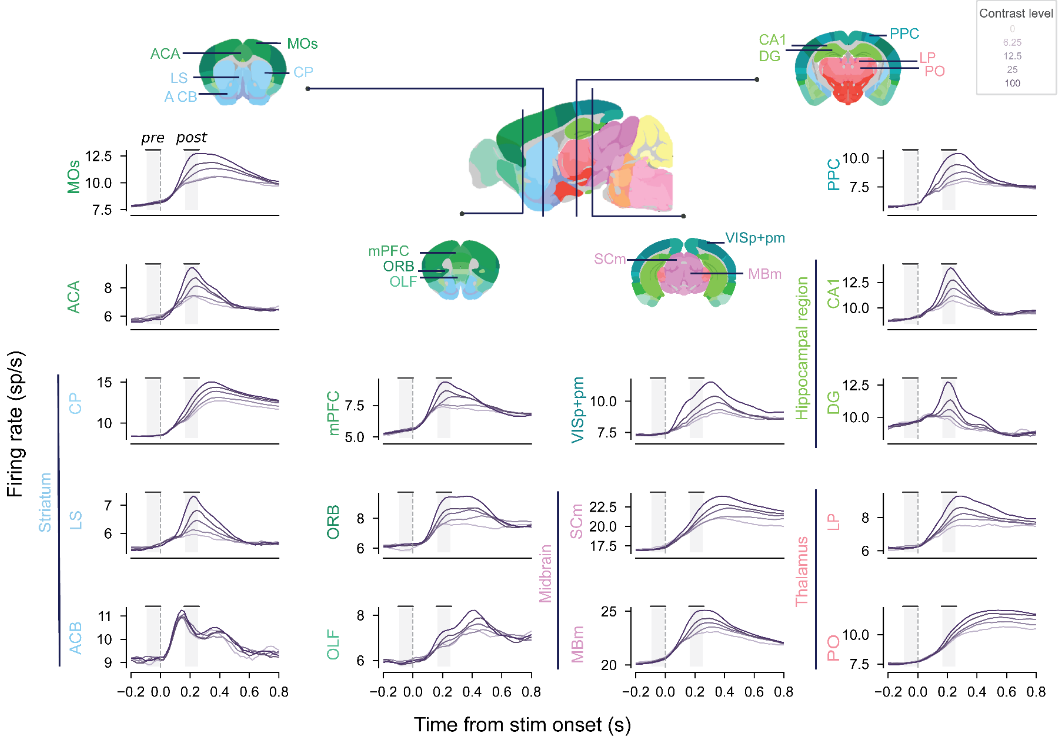


**Supplementary Figure 11, related to Figure 3. Regional specificity in contrast modulation of firing rate.** Time courses of the average firing rate in each ROI across different stimulus contrast levels, combining data from both age groups. Each line shows the mean firing rate across neurons for one contrast level. ROI-specific sample sizes are the same as in Supplementary Figure 10a and indicate the number of neurons. Grey shaded regions indicate the pre- and post-stimulus windows used for summary analyses.

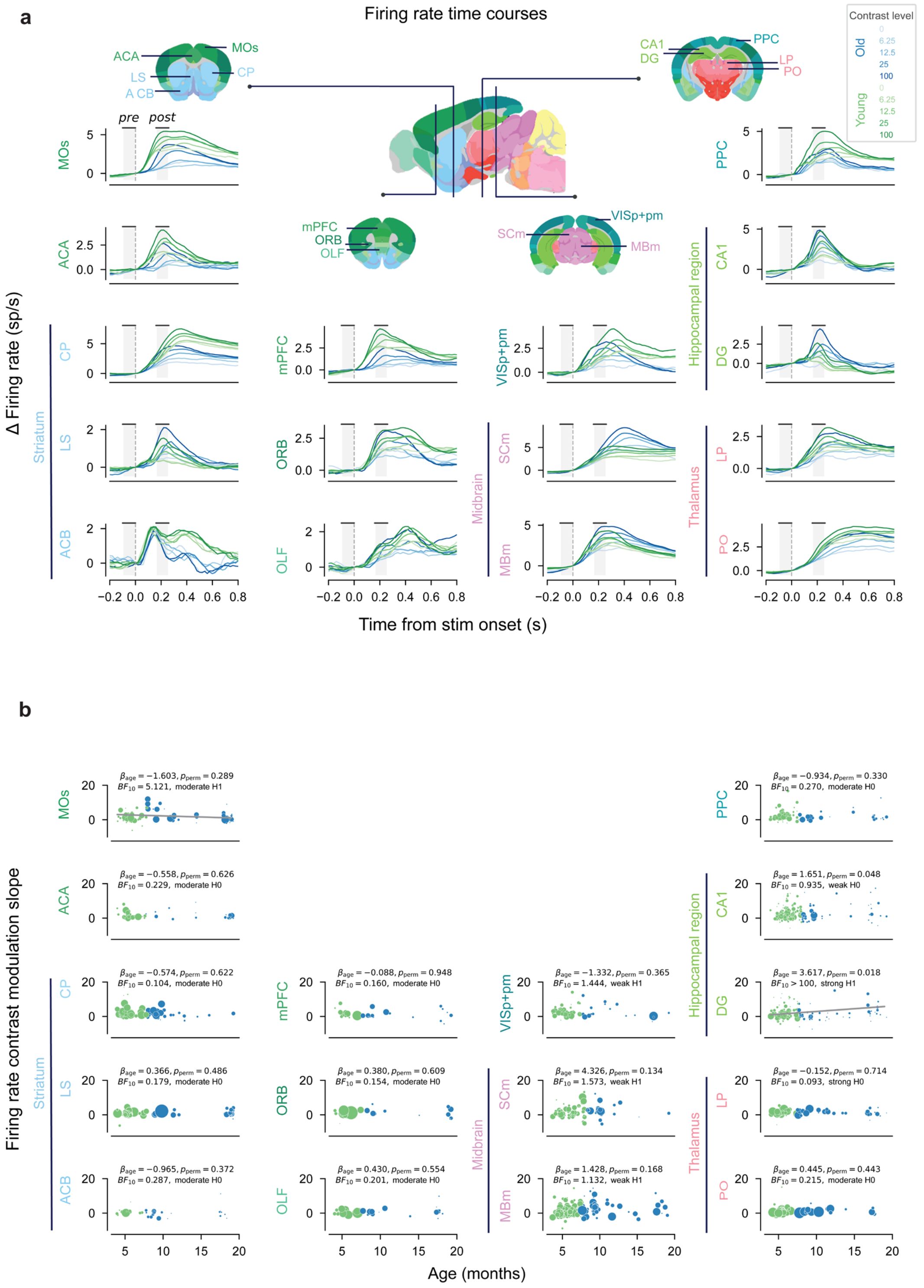


**Supplementary Figure 12, related to Figure 3. Regional specificity of age-related differences in contrast modulation of firing rates. (a)** Baseline-corrected mean firing rate time courses for different stimulus contrast levels in each ROI, aligned to stimulus onset. Saturation represents different contrast levels. Lines show mean firing rates across neurons. **(b)** The relationship between contrast modulation slope and mouse age. Each dot represents one insertion. For visualization, ach dot represents the mean contrast modulation slope across neurons from one insertion, and dot size represents the number of neurons recorded in that insertion. ROI-specific sample sizes are the same as in Supplementary Figure 10a and indicate the number of neurons.

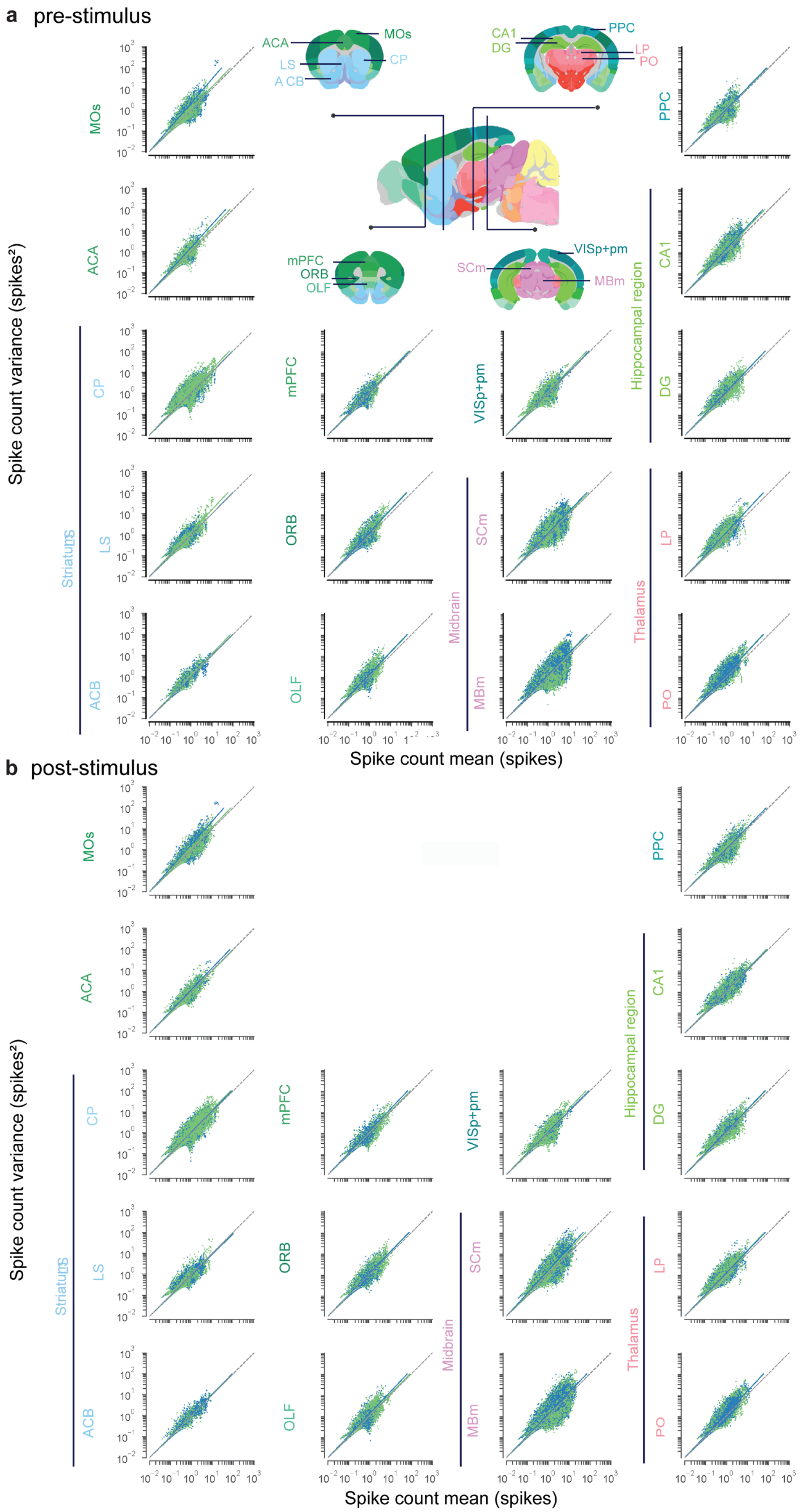
a pre-stimulus
ACA
MOs
LS
CP
A CB
CA1
DG
PPC
LP
PO
mPFC
ORB
OLF
SCm
VISp+pm
MBm
Spike count variance (spikes²)
Striatum
Hippocampal region
Midbrain
Thalamus
ACB
Spike count mean (spikes)
b post-stimulus
Spike count variance (spikes²)
Striatum
Hippocampal region
Midbrain
Thalamus
Spike count mean (spikes)

**Supplementary Figure 13, related to Figure 4. The relationship between spike count variance and spike count mean.** For each neuron and each stimulus level, we plotted its spike count variance (spikes$^2$) against its spike count mean (spikes) across trials, separately for (a) the pre-stimulus time window (-100ms to 0ms) and (b) the post-stimulus time window (160ms to 260ms). Colored lines represent linear fits for each age group (blue = old, green = young), with the slope corresponding to an estimate of the Fano Factor. We observed curvilinear patterns in the lower-left part of many panels, resembling patterns found in Figure 2 of a previous study[3]. The origin of these patterns remains unclear and warrants further investigation.

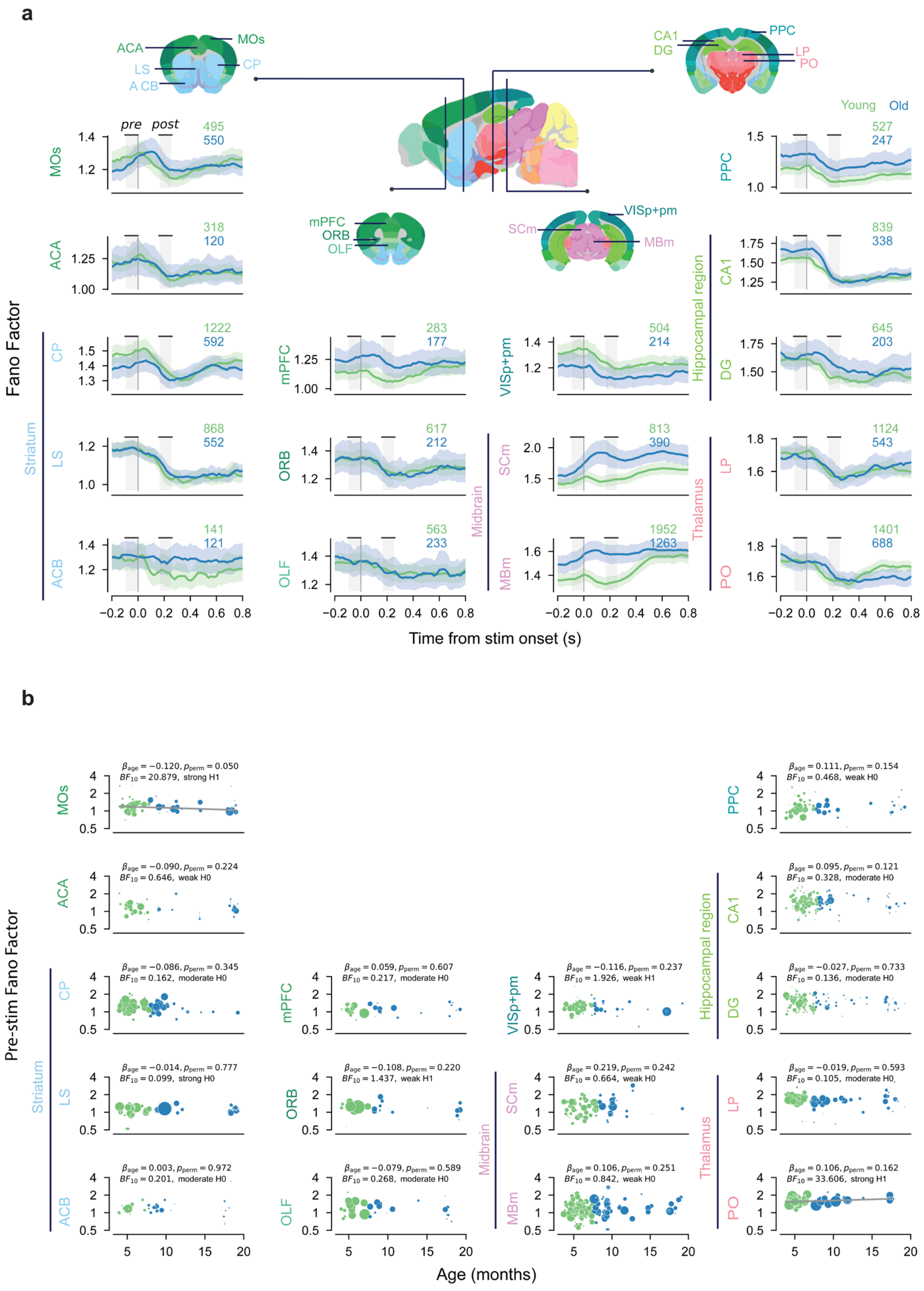

**c**

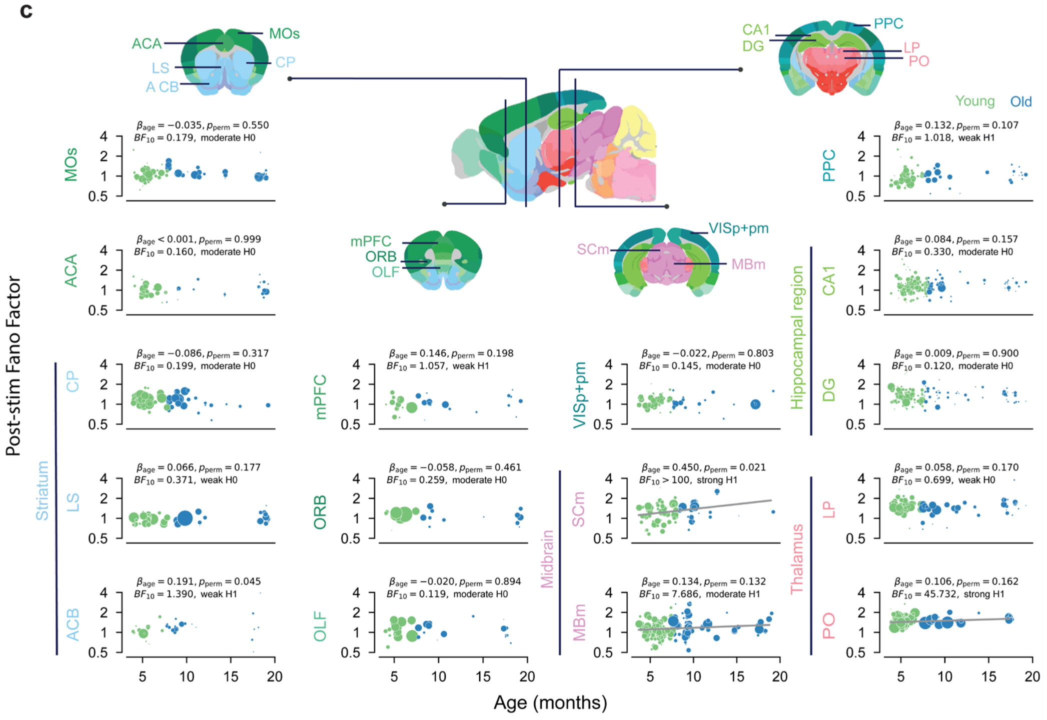


**Supplementary Figure 14, related to Figure 4. Regional specificity of mean-subtracted Fano Factors. (a)** Time courses of mean-subtracted Fano Factor in each ROI, aligned to stimulus onset. Thick lines show the mean Fano Factor across all neurons within each age group, with shaded areas representing the 95% confidence intervals. Sliding window width: 0.1 s; step size: 0.02 s. Numbers indicate the number of neurons in each group. **(b)** The relationship between pre-stimulus mean-subtracted Fano Factor and age. For visualization, each dot represents the mean across neurons from one insertion, and dot size represents the number of neurons recorded in that insertion. ROI-specific n values are the same as in (a) and indicate the number of neurons included in the analysis. **(c)**The relationship between post-stimulus mean-subtracted Fano Factor and age. Visualization, sample-size definition, plotting, and statistical conventions are the same as in (b).

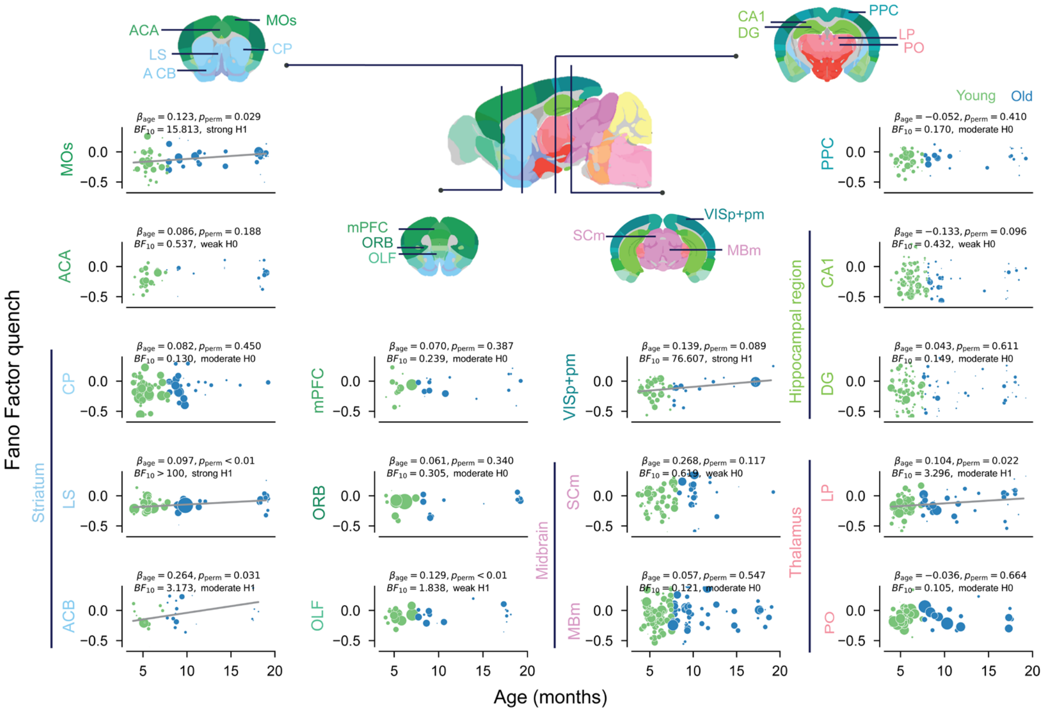


**Supplementary Figure 15, related to Figure 4. Regional specificity of mean-subtracted Fano Factor quenching.** The relationship between the mean-subtracted Fano Factor quench index (calculated as post-stimulus Fano Factor minus pre-stimulus Fano Factor for each neuron) and mouse age. For visualization, each dot represents the mean quench index across neurons from one insertion, and dot size represents the number of neurons recorded in that insertion. ROI-specific n values are the same as in Supplementary Figure 14a and indicate the number of neurons included in the analysis.

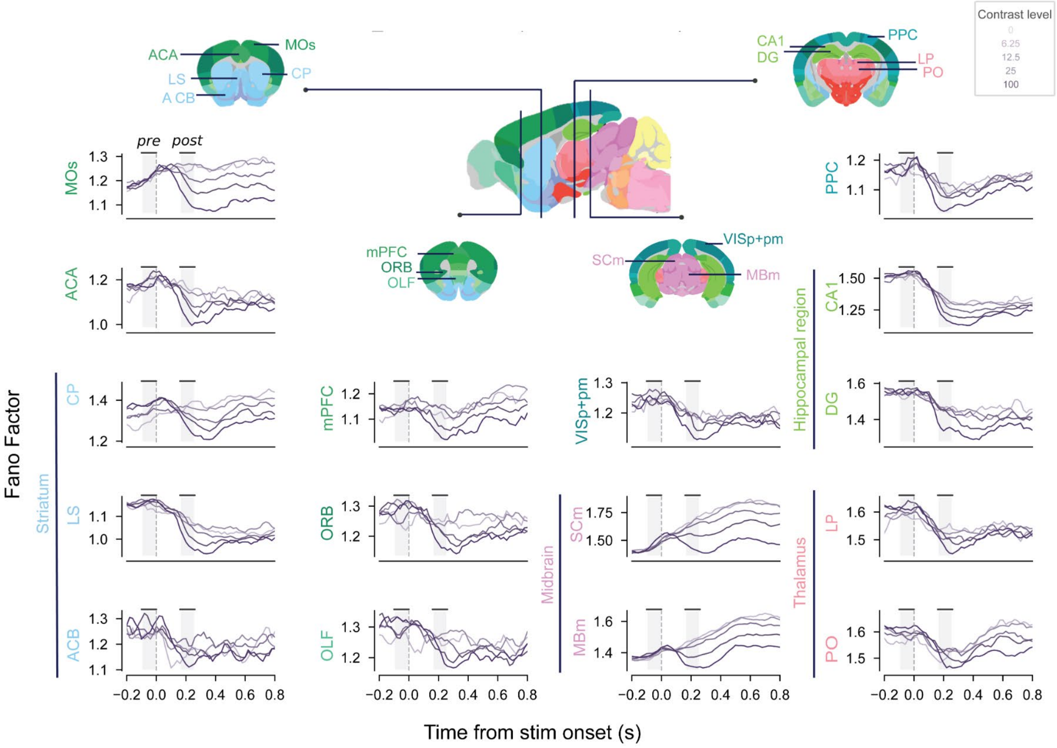

**Supplementary Figure 16, related to Figure 4. Regional specificity in contrast modulation of the Fano Factor.** Time courses of the mean Fano Factor for different stimulus contrast levels and ROIs, combining data from both age groups. Each line shows the mean Fano Factor across neurons for one contrast level. ROI-specific n values are the same as in Supplementary Figure 14a and indicate the number of neurons. Grey shaded regions indicate the pre- and post-stimulus windows used for summary analyses.

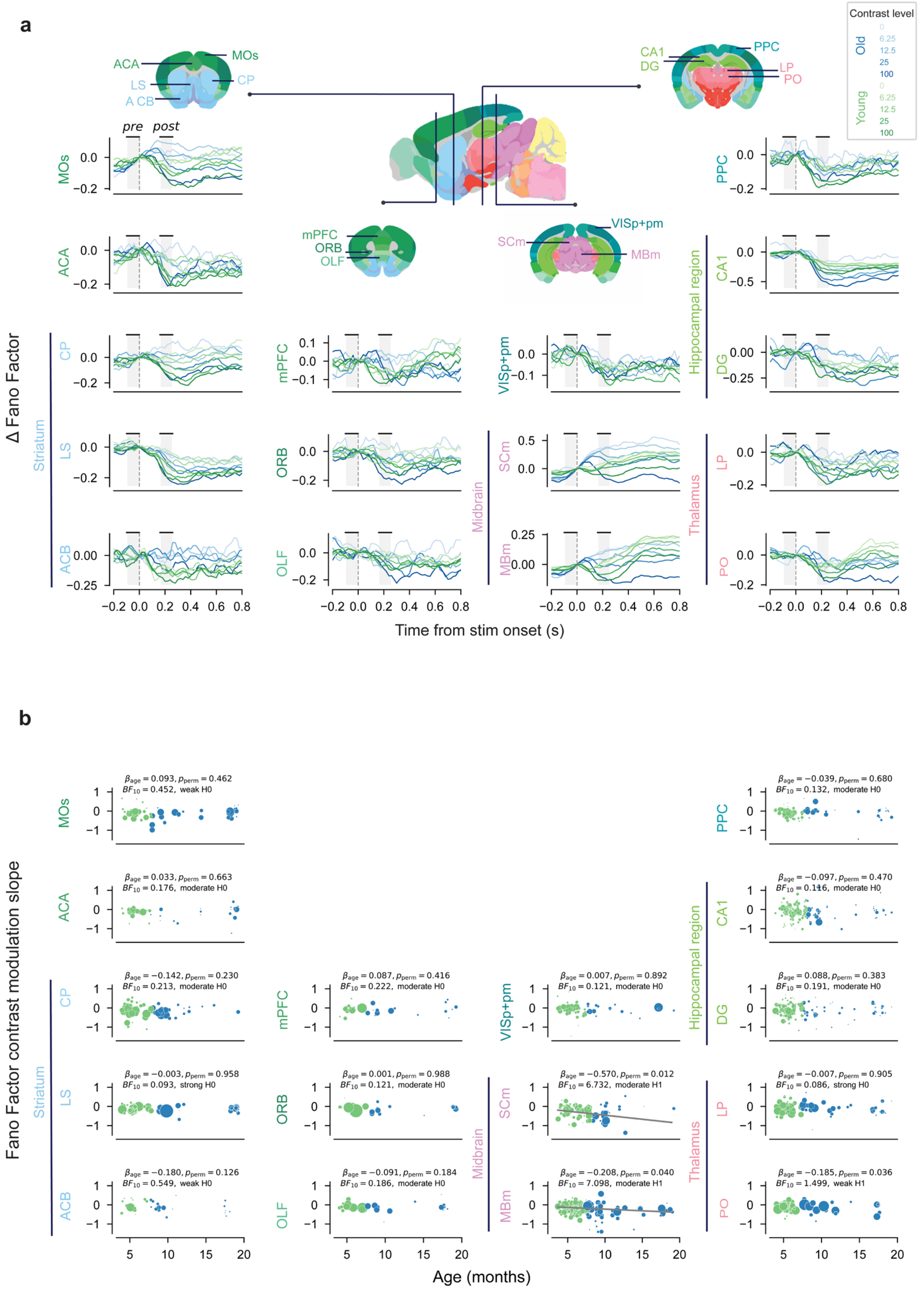


**Supplementary Figure 17, related to Figure 4. Regional specificity in age-related differences in contrast modulation of the Fano Factor. (a)** Baseline-corrected Fano Factor time courses, across neurons, for different stimulus contrast levels in each ROI, aligned to stimulus onset. Colors represent the age groups (blue = old, green = young). Saturation represents different contrast levels. Sliding window width: 0.1 s; step size: 0.02 s. Grey areas indicate the pre-stimulus (-100 to 0 ms) and post-stimulus (160 to 260 ms) time windows used for summary analyses. **(b)** The relationship between Fano Factor contrast-modulation slope and mouse age. Contrast modulation slopes were computed for each neuron by fitting a linear relationship between the change in baseline-corrected Fano Factor and the different stimulus contrast levels. For visualization, each dot represents the mean contrast-modulation slope across neurons from one insertion. ROI-specific n values are the same as in Supplementary Figure 14a and indicate the number of neurons used in the statistical analysis.

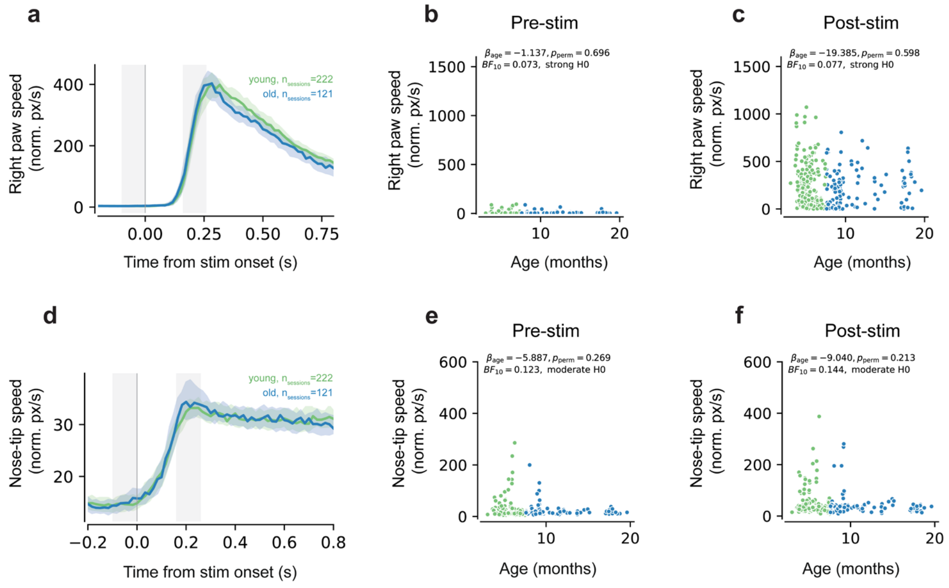


**Supplementary Figure 18, related to Figure 4. Right-camera DLC movement time courses and session-level summary metrics.** All panels use session-level data: n = 343 sessions (222 young and 121 old sessions). **(a)** Right paw speed time course aligned to stimulus onset (t = 0), summarized across sessions (median ± 95% confidence interval). Gray shaded regions indicate the pre-stimulus (−100 to 0 ms) and post-stimulus (160 to 260 ms) windows used to compute session-level movement summaries. **(b–c)** Session-wise median right paw speed within the pre-stimulus window (b) and post-stimulus window (c), plotted against mouse age; each dot corresponds to one session. **(d)** Nose-tip speed time course aligned to stimulus onset, summarized across sessions (median ± 95% confidence interval), with the same pre/post windows indicated. **(e–f)** Session-wise median nose-tip speed within the pre-stimulus (e) and post-stimulus (f) windows versus age; each dot corresponds to one session.

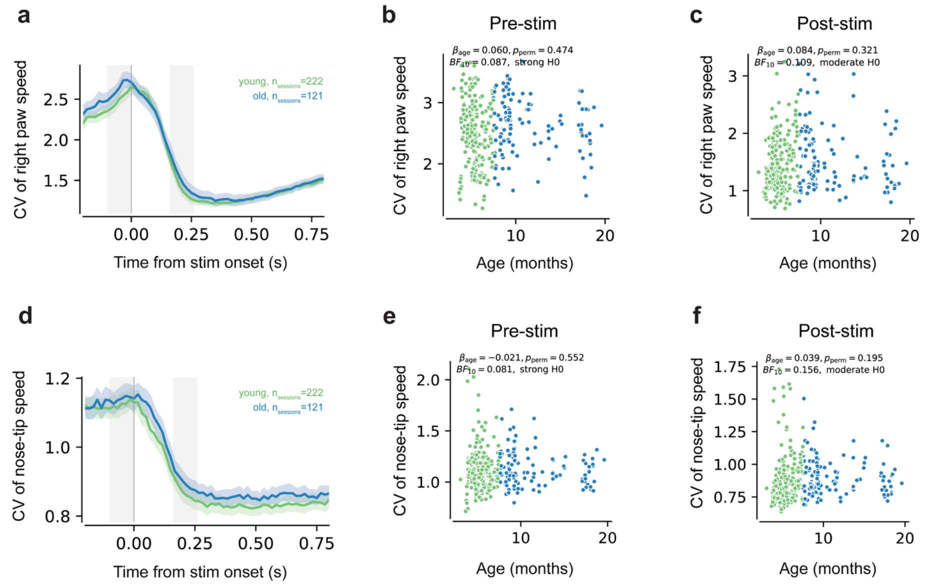


**Supplementary Figure 19, related to Figure 4. Right-camera DLC movement variability time courses and session-level summary metrics.** All panels use session-level data: n = 343 sessions (222 young and 121 old sessions). **(a)** Time course of right paw speed variability, quantified as the coefficient of variation (CV) across trials and aligned to stimulus onset (t = 0), summarized across sessions (mean ± 95% confidence interval). Gray shaded regions indicate the pre-stimulus (−100 to 0 ms) and post-stimulus (160 to 260 ms) windows used to compute session-level movement variability summaries. **(b–c)** Session-wise right paw speed CV within the pre-stimulus window (b) and post-stimulus window (c), plotted against mouse age; each dot corresponds to one session. **(d)** Time course of nose-tip speed variability, quantified as CV and aligned to stimulus onset, summarized across sessions (mean ± 95% confidence interval), with the same pre/post windows indicated. **(e–f)** Session-wise nose-tip speed CV within the pre-stimulus (e) and post-stimulus (f) windows versus age; each dot corresponds to one session.

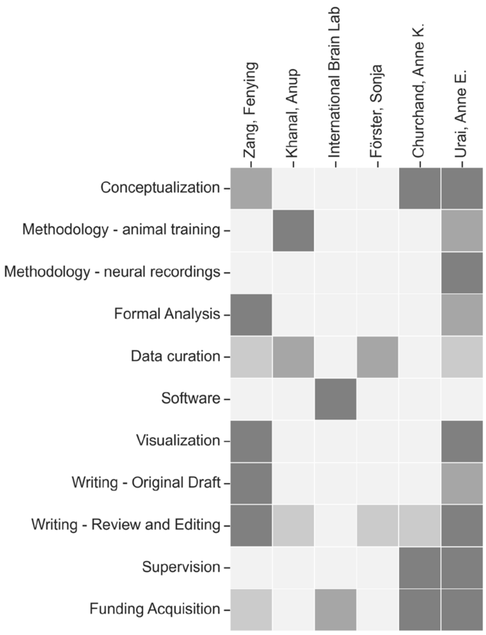


**Supplementary Figure 20. CRediT matrix of author contributions.**

### Supplementary Tables

| metric | β_age | BF_evidence_age |
|---|---|---|
| rt_median | -0.014 | <0.1, strong H0 |
| rt_CV | +0.158 | >100, strong H1 |
| pre_fr | +0.216 | >100, strong H1 |
| post_fr | +0.229 | >100, strong H1 |
| fr_delta_modulation | +0.373 | 0.28, moderate H0 |
| pre_ff | -0.001 | <0.1, strong H0 |
| post_ff | +0.056 | >100, strong H1 |
| ff_quench | +0.070 | >100, strong H1 |
| ff_quench_modulation | -0.066 | 1.42, weak H1 |

**Supplementary Table 1. Age effects from the original models (in the main analyses).** For each of the key behavioral and neural metrics, the table reports β and $BF_{10}$ for age.

| metric | β_age | BF_evidence_age | β_training | BF_evidence_training |
|---|---|---|---|---|
| rt_median | -0.025 | 0.22, moderate H0 | +0.483 | 0.49, weak H0 |
| rt_CV | +0.140 | >100, strong H1 | +0.276 | 0.37, weak H0 |
| pre_fr | +0.211 | >100, strong H1 | +0.311 | <0.1, strong H0 |
| post_fr | +0.202 | >100, strong H1 | +1.200 | >100, strong H1 |
| fr_delta_modulation | +0.394 | 0.42, weak H0 | -0.946 | <0.1, strong H0 |
| pre_ff | -0.012 | <0.1, strong H0 | +0.449 | 7.81, moderate H1 |
| post_ff | +0.045 | 13.6, strong H1 | +0.424 | 8.87, moderate H1 |
| ff_quench | +0.077 | >100, strong H1 | -0.297 | 0.25, moderate H0 |
| ff_quench_modulation | -0.072 | 3.16, moderate H1 | +0.310 | <0.1, strong H0 |

**Supplementary Table 2. Controls for training duration: extended models with age and training duration.** For each behavioural and neural metric, we report results on an extended model including both age at recording and training duration. In the extended models, the age term reflects the partial age effect controlling for training duration; the training term reflects any additional contribution of training duration beyond age.

| metric | β_age | BF_evidence_age | β_paw | BF_evidence_paw | β_nose | BF_evidence_nose |
|---|---|---|---|---|---|---|
| pre_fr | +0.201 | >100, strong H1 | +0.016 | 0.37, weak H0 | -0.028 | >100, strong H1 |
| post_fr | +0.211 | >100, strong H1 | +0.006 | <0.1, strong H0 | -0.027 | >100, strong H1 |
| fr_delta_modulation | +0.437 | 1.12, weak H1 | -0.086 | <0.1, strong H0 | -0.119 | 3.7, moderate H1 |
| pre_ff | +0.017 | <0.1, strong H0 | -0.015 | 6.14, moderate H1 | +0.006 | 0.11, moderate H0 |
| post_ff | +0.062 | >100, strong H1 | -0.014 | 7.58, moderate H1 | -0.003 | <0.1, strong H0 |
| ff_quench | +0.052 | 11.7, strong H1 | -0.002 | <0.1, strong H0 | -0.005 | >100, strong H1 |
| ff_quench_modulation | -0.076 | 5.81, moderate H1 | -0.012 | <0.1, strong H0 | +0.014 | 0.45, weak H0 |

**Supplementary Table 3. Additional controls using video-based movement (right camera): Extended model summary (Age + Paw + Nose-tip).** In sessions with QC-passing right-camera recordings (n = 343), we extracted right-paw speed (task-relevant) and nose-tip speed (less task-specific) in pre/post windows and added them as covariates in the extended models. The table reports β and $BF_{10}$ for age, paw, and nose-tip terms.

| metric | β_age | BF_evidence_age | β_paw_cv | BF10_paw_cv | β_nose_cv | BF10_nose_cv |
|---|---|---|---|---|---|---|
| pre_fr | +0.200 | >100, strong H1 | +0.020 | 3.07, moderate H1 | -0.017 | 1.11, weak H1 |
| post_fr | +0.207 | >100, strong H1 | +0.026 | 13.3, strong H1 | +0.023 | 4.83, moderate H1 |
| fr_delta_modulation | +0.552 | 6.59, moderate H1 | +0.130 | 0.77, weak H0 | -0.017 | <0.1, strong H0 |
| pre_ff | +0.014 | <0.1, strong H0 | +0.025 | >100, strong H1 | -0.009 | 0.28, moderate H0 |
| post_ff | +0.066 | >100, strong H1 | +0.023 | >100, strong H1 | -0.010 | 0.42, weak H0 |
| ff_quench | +0.046 | 2.89, weak H1 | +0.006 | >100, strong H1 | -0.007 | 1.55, weak H1 |
| ff_quench_modulation | -0.081 | 10.8, strong H1 | -0.000 | <0.1, strong H0 | +0.007 | <0.1, strong H0 |

**Supplementary Table 4. Additional controls using video-based movement variability (right camera): Extended model summary (Age + Paw CV+ Nose-tip CV).** In sessions with QC-passing right-camera recordings (n = 343), we extracted right-paw and nose-tip speed variability, quantified as CV within the relevant pre- and/or post-stimulus windows, and added these as covariates in the extended models. The table reports β and $BF_{10}$ for age, paw CV, and nose-tip CV terms. Although these covariates explained additional variance in several cases, adding these movement-variability measures as covariates to the extended models left the main age-related pattern broadly unchanged. For firing-rate contrast modulation, the evidence shifted from moderate evidence for H0 to weak evidence for H1 after including movement variability covariates, yielding weak evidence for an age effect in the adjusted model. For two additional metrics, the qualitative conclusion remained the same, but the strength of evidence changed: for Fano factor quenching, evidence for H1 was attenuated from strong to weak, whereas for Fano factor contrast modulation, evidence for H1 increased from weak to strong. All other metrics retained the same qualitative conclusion.
